# Mapping the biogenesis of forward programmed megakaryocytes from induced pluripotent stem cells

**DOI:** 10.1101/2021.04.21.440767

**Authors:** Moyra Lawrence, Arash Shahsavari, Susanne Bornelöv, Thomas Moreau, Katarzyna Kania, Maike Paramor, Rebecca McDonald, James Baye, Marion Perrin, Maike Steindel, Paula Jimenez-Gomez, Christopher Penfold, Irina Mohorianu, Cedric Ghevaert

## Abstract

Platelet deficiency, known as thrombocytopenia, can cause haemorrhage and is treated with platelet transfusions. We developed a system for the production of platelet precursor cells, megakaryocytes, from pluripotent stem cells. These cultures can be maintained for >100 days, implying culture renewal by megakaryocyte progenitors (MKPs). However, it is unclear whether the MKP state *in vitro* mirrors the state *in vivo*, and MKPs cannot be purified using conventional surface markers. We performed single cell RNA sequencing throughout *in vitro* differentiation and mapped each state to its equivalent *in vivo*. This enabled the identification of 5 surface markers which reproducibly purify MKPs, allowing us an insight into their transcriptional and epigenetic profiles. Finally, we performed culture optimisation, increasing MKP production. Altogether, this study has mapped parallels between the MKP states *in vivo* and *in vitro* and allowed the purification of MKPs, accelerating the progress of *in vitro*-derived transfusion products towards the clinic.

## Introduction

Platelets are the cells responsible for clotting. Every year in the UK, 280,000 platelet units are transfused into patients with a low platelet count, known as thrombocytopoenia, to prevent haemorrhage (Heckman et al., 1997). The majority of patients receive platelet transfusions either after haemorrhage or bone marrow insufficiency following cancer treatment or haematopoietic malignancy (Estcourt et al., 2017, Slichter, 1980, Harker and Finch, 1969). Platelet units are collected from donors (Körbling and Freireich, 2011, Stroncek and Rebulla, 2007) and matched for ABO Blood Type and Rhesus D (Estcourt et al., 2017). However, previous transfusion recipients and multiparous women may become immunised against foreign MHC Class-I or other antigens on the platelet surface, adding even more complexity to the matching process (Kickler et al., 1990, Kiefel et al., 2001, Stanworth et al., 2015). In addition, short platelet shelf-life (5-7 days) and restricted donor availability mean that platelets have the most precarious supply chain of all blood components and shortages may occur.

Megakaryocytes (MKs) are large, multinucleated, polyploid cells (Yamada, 1957, Sola-Visner et al., 2007) which differentiate from haematopoietic stem cells (HSCs) in the bone marrow (Machlus and Italiano, 2013). MKs make up 0.01-0.03% of nucleated bone marrow cells (Nakeff and Maat, 1974) and collectively they are estimated to produce 1-2 x 10^11^ platelets daily (Bluteau et al., 2009, Harker, 1977). The long-standing model of HSC differentiation is a hierarchical model (Figure 1a) where HSCs differentiate to form multipotent progenitors (MPPs) which undergo transit-amplifying divisions and can generate all mature blood cells (Morrison and Weissman, 1994). As these differentiate and become more lineage restricted along the path to MK differentiation, common myeloid progenitors (CMPs) are formed (Deutsch and Tomer, 2006, Manz et al., 2002, Akashi et al., 2000) that subsequently differentiate to become megakaryocyte erythroid progenitors (MEPs), which are restricted to MK and erythrocyte production (Deutsch and Tomer, 2006) influenced by *MYB* (Sanada et al., 2016), *KLF1*/*FLI1* (Chen et al., 2014) and the cell cycle (Lu et al., 2018). Reduced *MYC* is required for MK specification (Lu et al., 2018). MK progenitors (MKPs) differentiate from MEPs and can only form MKs (Deutsch and Tomer, 2006, Psaila et al., 2016). These were first observed in the human MEPs produced from mobilised peripheral blood-derived CD34^+^ cells (Psaila et al., 2016). MKPs retain very little capacity to produce erythroid cells and almost exclusively produce MKs in colony forming assays (Psaila et al., 2016). They express MK-associated proteins such as CD41, CD61, VWF, CLU and NF1B with CD42 expression heralding the extinction of erythroid colony forming potential (Psaila et al., 2016).

**Figure 1:**
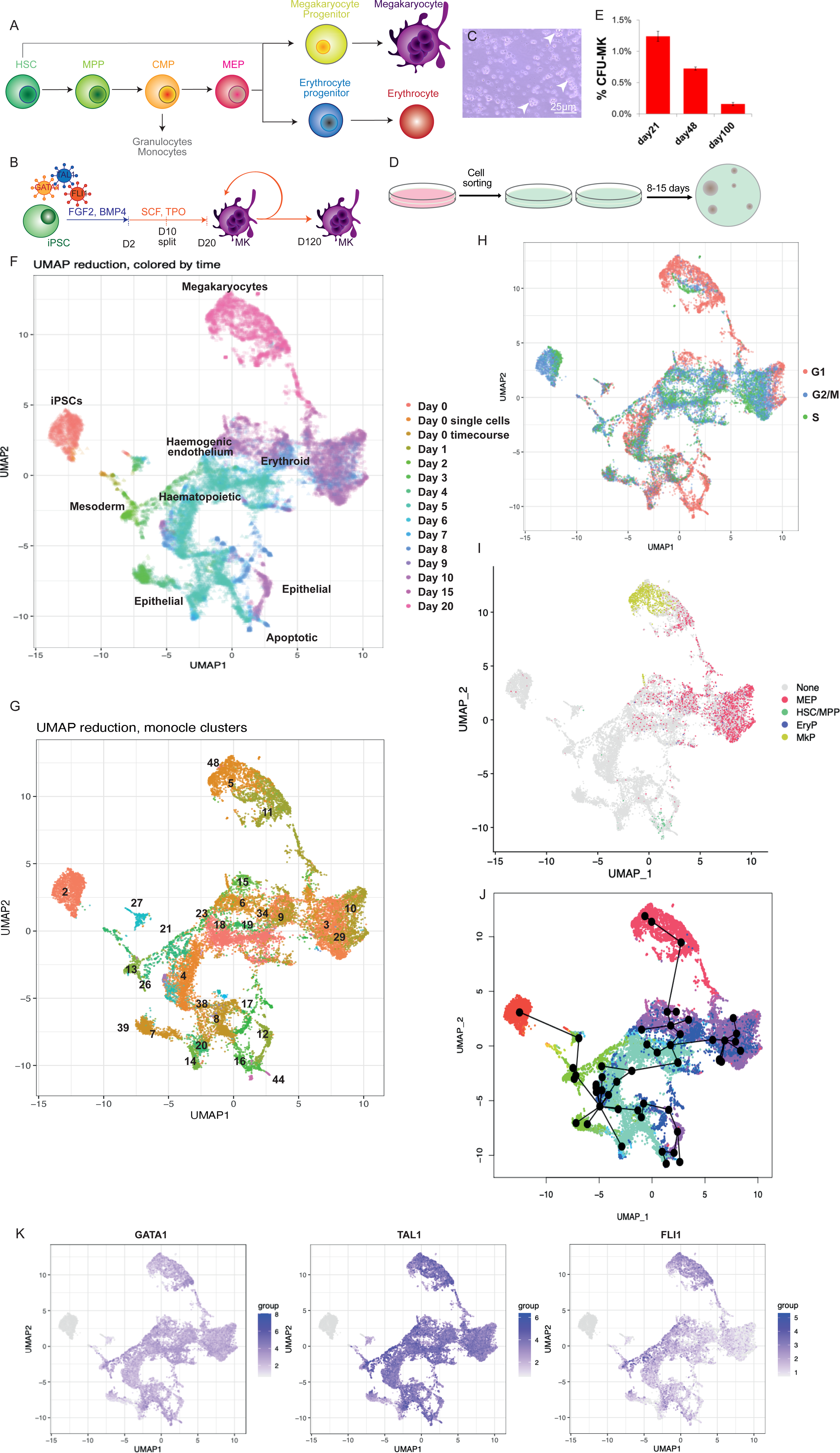
Single cell sequencing of MK differentiation from Induced Pluripotent Stem Cells allows the comparison to human *in vivo* haematopoiesis and trajectory analysis. **A.** Overview of haematopoiesis *in vivo*. HSC: Haematopoietic stem cell, MPP: multipotent progenitor, CMP: Common Myeloid Progenitor, MEP: Megakaryocyte Erythroid progenitor. **B.** Schematic of protocol (Moreau et al., 2016). iPSCs are lentivirally transduced with *GATA1*, *TAL1* and *FLI1*. After 20 days, mature megakaryocytes (MK) are formed which can persist for up to 120 days. **C.** Brightfield image of A1ATD1 MKs on D20 of differentiation. Arrowheads: proliferative clusters. Scale bar 25μm. **D.** Schematic of colony forming unit (CFU) assay. MKs are sorted into semi-solid methylcellulose containing SCF and TPO. After 9-15 days, emergent colonies can be scored. **E.** Percentage colony formation in an A1ATD1 culture in CFU assays at D21, D48 and D100 of culture. **F.** UMAP embedding of single cell RNA sequencing of MK differentiation time-course (10x Genomics). Legend: timepoint during differentiation. **G.** Monocle clusters on time-course data; 51 clusters identified using density peak-based clustering. Clusters mentioned in the text are labelled. Full plot Figure S3a. **H.** UMAP of cell cycle stage, inferred from the expression of cell cycle-associated transcripts. **I.** Random forest-predicted cell types on single cell time-course data. Cell types learned on 10X haematopoietic stem and progenitor cells from human bone marrow, spleen and peripheral blood. MEP: Megakaryocyte Erythroid Progenitor, HSC/MPP: Haematopoietic Stem Cell/Multipotent Progenitor. MkP: Megakaryocyte Progenitor, EryP: Erythroid Progenitor. H: Donor 1, 1% D0 cutoff, threshold of 0.53. **J.** Slingshot trajectory analysis of clusters from G coloured by timepoint. **K.** UMAP of un-normalised *GATA1*, *TAL1* and *FLI1* expression from lentiviral transgenes.

There is also evidence to suggest that MKs can differentiate directly from earlier haematopoietic intermediates. Distinct subsets of HSCs/MPPs express MK markers while still in the stem cell compartment (Notta et al., 2016, Wilson et al., 2015, Novershtern et al., 2011) from which MKs may be able to differentiate directly in times of stress (Notta et al., 2016). MKPs can also emerge directly from CMPs without becoming MEPs (Miyawaki et al., 2017).

After the discovery of induced pluripotent stem cells (iPSCs) they were rapidly adopted as a source for *in vitro* platelet generation as they were easily expandable, unlike HSCs. Protocols for MK production from iPSCs can be divided into two main groups: one relying on sequential cytokine cocktails to guide PSCs through developmental, termed directed differentiation (Feng et al., 2014, Mills et al., 2014, Börger et al., 2016, Takayama et al., 2008, Eicke et al., 2018, Lu et al., 2011, Gaur et al., 2006, Pick et al., 2013) and the other involving the overexpression of key transcription factors, termed forward programming (Moreau et al., 2016, Elcheva et al., 2014). The latter tends to produce higher purity MKs and use minimal cytokines. Some groups have also produced immortalised MKPs which makes the culture easily scalable but may have safety drawbacks due to genetic instability and a higher tumorigenicity risk to recipients (Nakamura et al., 2014, Takayama and Eto, 2012, Suzuki et al., 2020, Ito et al., 2018). iPSCs can be genetically modified (Zou et al., 2009), promoting efficient differentiation using inserted cassettes for the overexpression of differentiation-promoting factors (Pawlowski et al., 2017, Evans et al., 2021) and allowing MHC Class-I deletion to reduce the immunogenicity of manufactured platelets (Feng et al., 2014, Suzuki et al., 2020).

We previously described a protocol for mature MK differentiation from iPSCs using two cytokine combinations (Moreau et al., 2016) through the overexpression of *GATA1*, *TAL1* and *FLI1*. These MKs produce functional platelets which can contribute to thrombus formation (Shepherd et al., 2018). Interestingly, early timepoints of the culture produce both erythroid cells and MKs, tuned by the rheostat of *FLI1* expression (Dalby et al., 2018), mirroring *in vivo* megakaryopoiesis (Chen et al., 2014). One of the key characteristics of the cultures is their ability, beyond reaching maturity at day 20 (D20), to survive for a further 100 days whilst maintaining cell proliferation and purity by CD41 and CD42 expression (Moreau et al., 2016), indicating the presence of cells capable of reconstituting the culture when mature MKs fragment to generate platelets. We hypothesised that the culture is maintained by highly proliferative MKPs, mirroring MKPs *in vivo*.

Here we used single cell RNA sequencing to map differentiation from iPSCs to mature MKs *in vitro.* We investigated whether the progenitors generated during *in vitro* MK differentiation mirrored haematopoietic progenitors *in vivo* and found that only the more mature MKs and erythroid cells had a transcriptional signature similar to human primary MEPs and MKPs. Using a combination of single cell RNA sequencing and machine learning approaches on index sort data, we identified a combination of 5 novel surface markers that reproducibly isolated and purified MKPs from bulk cultures. MKPs purified using this approach provided new insights into MKPs’ transcriptional and epigenetic state. Finally, we optimised culture conditions for the expansion of MKPs and showed that this increased culture efficiency and postponed long-term culture exhaustion, potentially facilitating subsequent large-scale manufacturing of a clinical product.

## Results

### Forward programmed MK cultures from iPSCs contain MK progenitors

We previously described a robust and reproducible protocol for the production of mature, functional megakaryocytes (MKs) from induced pluripotent stem cells (iPSCs), catalysed by the lentiviral overexpression of *GATA1*, *TAL1* and *FLI1* (Figure 1b). The culture is adherent until D10 when it is disrupted and put into suspension. Mature MKs are harvested from D20. The cells acquire CD235 expression, then CD41 expression and finally CD42 expression as they become mature (Dalby et al., 2018, Moreau et al., 2016) and the cultures can produce a constant supply of mature MKs for up to 120 days. Proliferating cell clusters are often seen in these cultures (Figure 1c), suggesting that they are maintained by dividing MK progenitors (MKPs) at the centre of each cluster. Progenitors can be quantified in colony formation assays whereby a suspension of cells is seeded into a semi-solid medium containing cytokines. Each progenitor will form a “colony” that can be counted using microscopy and phenotyped by flow cytometry. Colony assays (Figure 1d) performed on live-sorted MK cultures suggested that colony-forming MKPs make up only around 1% of early cultures and diminish as the culture ages (Figure 1e). This confirmed that as the progenitors disappear, the cultures become exhausted. Morphologically, these progenitors are identical to the surrounding cells and cannot be identified using classical haematopoietic and/or MK surface markers (such as CD34, KIT, KDR or CD61 (Figure S1A and S1B)).

### Mapping MK differentiation *in vitro* using single cell RNASeq

To understand the processes of MK differentiation *in vitro*, we fixed A1ATD1 cells for 10x Genomics scRNA-seq every 24h during the first 10 days of the MK differentiation protocol and also on D15 and D20 when the MKs are mature. We also designed a unique strategy for transgene fingerprinting (Figure S1c): the cells had been differentiated with the aid of three lentiviral transcription factors, stably inserted into the genome. Their downstream lentiviral sequences allowed the amplification of their transcripts from cDNA with specially designed primers which were substituted for the V(D)J primers during target enrichment. This allowed an overview of the gene expression of these progenitors and also the transgene quantities catalysing their formation.

Quality control was performed as detailed in Figure S2 and STAR methods. All cells were projected on a Uniform Manifold Approximation and Projection (UMAP), coloured by timepoint (Figure 1f, Figure S2m). Fifty-one clusters were detected by Seurat and we set Monocle to find the same number of clusters (Figure 1g and Figure S3a). The set of cluster markers and enrichment terms (Supplementary Tables 3 and 4) were used to assess cluster identity and function.

iPSCs were located at the top left of the UMAP (Figure 1f) and separated into three clusters (“iPSC” cluster 2, 21 and 27) of which the main cluster (cluster 2) had the highest potency, expressing higher levels of markers of the naïve (Figure S3b) and primed pluripotent states (Figure S3c). As cells left cluster 2, they lost the expression of pluripotency-associated markers and acquired a mesoderm identity, expressing *HAND1* (“Mesoderm” clusters 4, 7, 8, 13, 26, 38, 39 and 43, Figure S3d). From here, the cells bifurcated onto two paths. The first progressed towards the bottom of the UMAP plot and expressed higher levels of mitochondrial transcripts. Some of these cells apoptosed at the bottom when the cells were dissociated from adhesive culture on D10 as seen by the enrichment of cluster 44 for apoptosis-associated GO terms. Other cells expressed epithelial markers such as *KRT8*, *KRT18* and *KRT19* (“Epithelial” clusters 8, 12, 14, 16, 17, 44, 45 and 47, Figure S3d) indicating that they moved towards an epithelial fate. These cells were lost as the cells are dissociated on D10. A second branch of cells progressed towards the central part of the graph and expressed markers classically associated with haematopoiesis and haemogenic endothelium such as *CD34*, *KDR*, *CDH5 and PECAM1* (“Haemogenic endothelium” clusters 4, 18, 19, 21 and 23, Figure S3d). *GATA1* and *TAL1*, haematopoietic transcription factors were also expressed at this stage (Figure S3d). From D9 the cells transitioned towards an erythroid identity on the top right, characterised by *GYPA*, heme and porphyrin synthesis and iron metabolism (“Erythroid” clusters 3, 6, 9, 10, 15, 29, 33, 34, 37, 46 and 50). As these cells matured and moved towards the top of the graph, they acquired platelet-associated transcripts such as *ITGA2B* and finally became mature MKs expressing *GP1BA* and high levels of *FLI1* (“Megakaryocytes” clusters 5, 11 and 48) (Figure S3d).

We would expect MKPs to be found from an early time point in the culture as MK-producing cells are present before D10 (Dalby et al., 2018). MKPs should also persist until the end of the time-course as they support long-term culture. MKPs should therefore be found in clusters containing cells from multiple timepoints such as the “Haemogenic endothelium” clusters or the “Erythroid” clusters, both of which contained cells from D8 to the end of the time-course.

The progenitors we observe in colony forming assays are highly proliferative. Thus we carried out cell cycle mapping of the differentiation process, partitioning cells into stages of the cell cycle according to their dominant transcripts. Consistent with reports that human pluripotent stem cells have a shortened G1 phase (Becker et al., 2006, Ghule et al., 2011), the iPSCs on the top left had few cells in G1 (Figure 1h). Correspondingly, as the iPSCs began to differentiate, they entered G1 phase, consistent with reports of G1 lengthening during differentiation (Ruiz et al., 2011). The erythroid cluster and the MK cluster segregated based on cell cycle stage. Most of the MKs were in G1 phase, consistent with previous reports that G1-associated cyclins are required for the endomitosis involved in MK polyploidisation (Wang et al., 1995, Ravid et al., 2002). Interestingly, differentiating cells also progressed to G1 phase as they neared the MK state towards the top right of the plot, indicating that one of these G1-dominant clusters may contain MKPs.

The cell classifications used in Figure 1f were based on transcript enrichment relative to the rest of the dataset, which tells us that the MKs are, for example, more megakaryocytic than the rest of the dataset. It does not tell us, however, whether we generate cells which correspond to their *in vivo* counterparts. To investigate this, we compared our dataset to a dataset (Mende et al., 2020) containing 30,873 10x Genomics sequenced CD19^-^CD34^+^ human haematopoietic stem and progenitor cells (HSPCs) from the bone marrow, peripheral blood and spleen of two donors. A random forest model was trained on the cell types from the *in vivo* data, and deployed to predict the cell type identities of the time series data (further details available in the methods section). The results are shown in Figure 1i (donor 1) and Figure S3e (donor 2). Very few primary HSCs or MPPs mapped to the iPSC differentiation dataset, indicating that the forward programming of iPSCs to MKs is a process which specifies MK identity without passing through a HSC or HSC-like intermediate stage (Supplementary Table 5). From D9, when the cells began to express CD41, a marker of Megakaryocyte Erythroid Progenitors (MEPs), we saw cells emerging that mapped to primary MEPs. This is consistent with our previously published results demonstrating that D9-10 cells can undergo either MK or erythroid differentiation (Dalby et al., 2018). Most cells generated after D9 mapped directly to primary MKPs. Interestingly, on D9 the cells mapping to *in vivo* MKPs were the yellow points in cluster 6, making this cluster an attractive one for investigation as a potential MKP cluster (Figure 1i). We concluded from this data that forward programming enforces an accelerated patterning which bypasses the usual haematopoietic hierarchy but generates end stage progenitors which faithfully recapitulate primary MKPs.

To map the path of the cells through differentiation, we performed trajectory analysis through the monocle clusters using Slingshot (Street et al., 2018) (Figure 1j, Figure S4a). The trajectory began with markers such as *EPCAM*, which is highly expressed in human pluripotent stem cells (Sundberg et al., 2009, Kolle et al., 2009). The cells differentiated to mesoderm marked by transcripts such as *SMARCC*, which plays a key role in mesoderm formation during haematopoietic differentiation (Piazzi et al., 2015). Some of the cells became epithelial at the bottom of the graph, defined by transcripts associated with epithelial-to-mesenchymal transition such as *S100A4*. Other cells progressed towards early haematopoietic cells, propelled by key bone marrow niche-associated transcripts such as *RNPEP* (Nehme et al., 2015) and cytokinesis factors as *CLIC4* (Huang et al., 2020). Cells directly after this decision point (clusters 4, 20, 21) were characterised by the expression of ribosomal transcripts, which also characterised the end of the trajectory towards mature MKs, indicating increased protein synthesis. A crucial part of the trajectory was seen in clusters 18 and 19, where the trajectory divided into MK and erythroid fates. Markers upregulated here were also upregulated in the MK clusters and the erythroid clusters. These cells can form both mature MKs and erythroid cells (Dalby et al., 2018), demonstrating the functional relevance of the MEP state identified by trajectory analysis. The erythroid branch was defined by markers such as *TAL1*, which is associated with erythropoiesis (Pick et al., 2013). Finally, the branch of the trajectory corresponding to mature MKs was defined by transcripts such as *PF4* and *GP9* which are highly expressed in mature MKs (Thompson et al., 1996, Lanza, 2006). Trajectory analysis (Figure 1j) allows us to pinpoint clusters 6, 15 and 34 (Figure 1g) as clusters through which cells must pass to become mature MKs, implying the presence of MKPs in these clusters.

Next, we carried out targeted sequencing of lentiviral transcription factor-derived transcripts to investigate their contribution to specific trajectories (Figure 1k and Figure S4b). *GATA1* and *TAL1* were highly expressed throughout differentiation to all cell fates, however *FLI1* was expressed exclusively along the trajectory to mature MKs, with erythroid and epithelial clusters losing *FLI1* expression. This confirmed previous experimental results demonstrating that *FLI1* is instrumental in MK commitment and maturation (Dalby et al., 2018, Chen et al., 2014). Interestingly, clusters 6, 15 and 34 expressed more *FLI1* than other clusters from the same time-points, indicating that these clusters contained cells within the erythroid cluster which were destined to become MKs and could thus correspond to MKPs.

Together, our mapping allowed us to identify clusters 6, 15 and 34 (Figure 1g) as particularly interesting and potentially containing MKPs. These clusters contained cells from D8 onwards, as we would expect from an MKP cluster (Figure 1f). These clusters expressed more *FLI1* than other cells from the same time-points (Figure 1k). Finally, cells must pass through these clusters on their predicted trajectory towards the MK fate (Figure 1j). We therefore hypothesised that this region corresponded to MKPs.

### Identifying MK progenitor markers using a single cell approach

We hypothesised that MKPs have a more proliferative transcriptional signature than the surrounding cultures and must be found in mature MK cultures. To investigate this, we sequenced 192 single cells from a D40 A1ATD1(Yusa et al., 2011) MK culture (99% CD41^+^, 76% CD42^+^) using the Smart-Seq protocol. The increased sequencing depth allowed us to investigate possible cell-surface associated markers which could be used to purify the MKPs from mixed cultures.

After data filtering and analysis (STAR methods), three clusters were identified using Monocle, (Figure 2a). Transcript enrichment analysis allowed us to identify intermediate MKs which expressed *ITGA2B* at reduced levels (Group 1), mature MKs with high *ITGA2B* expression (Group 2) and apoptotic cells (Group 3). We found the first cluster most interesting due to their reduced maturity. We performed gene set enrichment analysis on the top 25 markers of each cluster (STAR methods). All 44 enriched terms enriched in cluster 1 were involved in DNA replication with E2F being the top enriched transcription factor (Supplementary Table 6). This indicated that this cluster is highly replicative and thus could contain MKPs.

**Figure 2:**
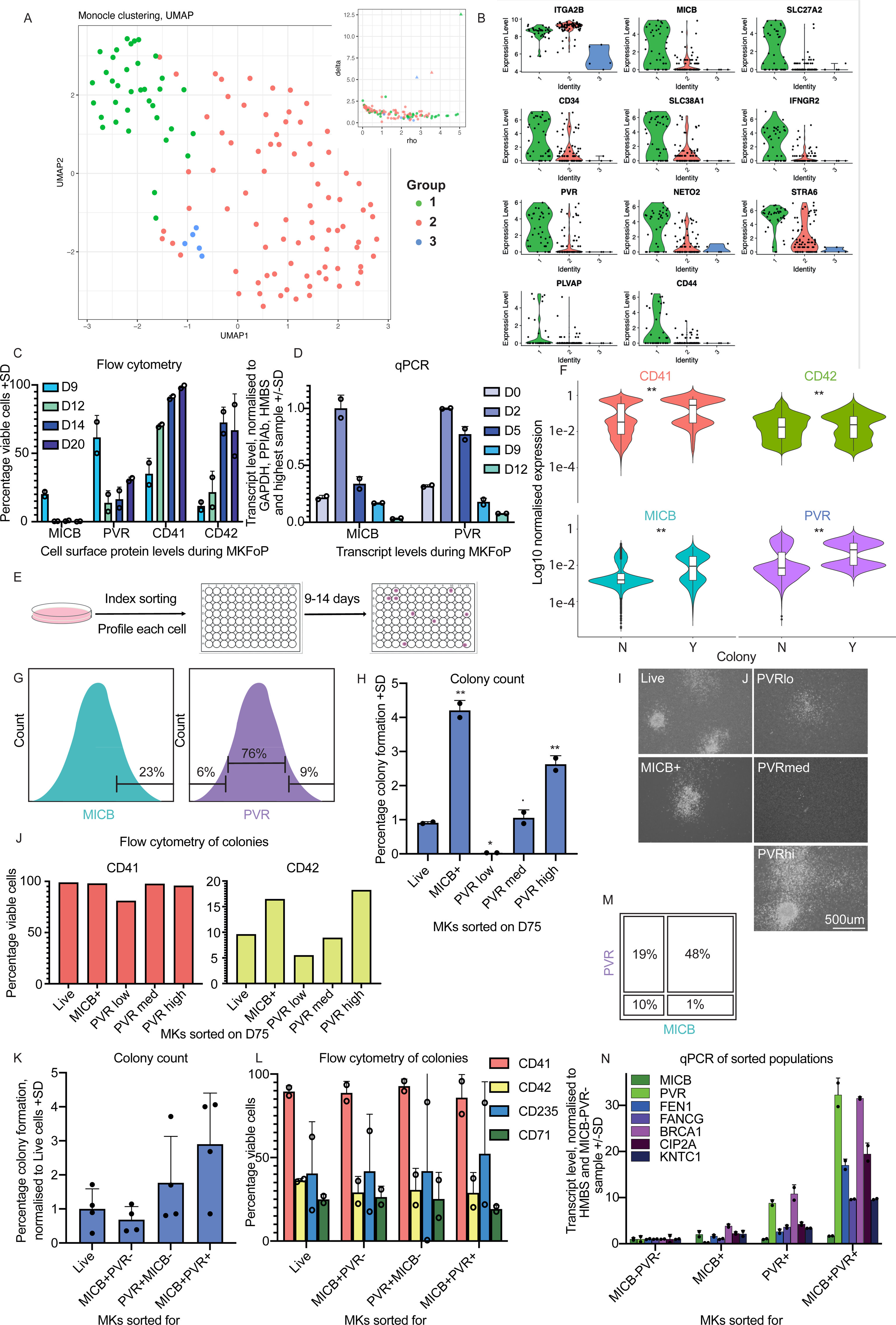
Identifying surface marker of megakaryocyte progenitor cells by single cell RNA sequencing. **A.** UMAP of SMART-Seq single cell RNA sequencing of a D44 A1ATD1 culture. Insert: density clustering decision plot. Delta (y-axis): local density of a cell. Rho (x-axis): distance to another cell with higher density. Triangles in upper right: cells with high values for rho and delta (cluster centres). **B.** Normalised expression levels of putative cell surface markers in each cluster from A. **C.** Flow cytometry analysis of MICB, PVR, CD41 and CD42 expression during differentiation of FFDK and QOLG3 iPSCs. **D.** qPCR analysis of *MICB* and *PVR* transcript levels during MK differentiation of A1ATD1 iPSCs. Transcript levels were normalised to the geometric mean of *GAPDH*, *PPIAb* and *HMBS* and the transcript level in the highest sample. **E.** Schematic of index sort. Single cells are stained and sorted into 96 well or 384 well plates containing medium supplemented with SCF and TPO. 9-14 days later, colonies are scored and colony-forming potential correlated to the cell surface marker expression of the parent cell. **F.** Results of index sorts of A1ATD1 MKs. Differences in expression between cells that did (Y) and did not form a colony (N) respectively measured by Wilcoxon test. CD41 (n=7), CD42 (n=7), MICB (n=6 with two different fluorophores), PVR (n=8). **G.** Sorting scheme: top 23% of MICB or high (top 9%), medium (middle 76%) or low (bottom 6%) of PVR. **H.** Colony counts on D12 after D75 A1ATD1 MKs were sorted for MICB or PVR. Colony counts were normalised to plated cell number and calculated as a percentage of input cells. Adjusted p value: compared to Live cells by one way ANOVA using Dunnett’s multiple comparisons test. **I.** Brightfield microscopy images of colonies in L. Scale bar 500μm. **J.** Flow cytometry analysis of cells from the colonies in L and parental MK culture. **K.** Colony counts on D11 or D13 after D33 or D35 A1ATD1 MKs respectively were sorted into CFU assays (top 50% of MICB, top 20% of PVR). Colony counts were normalised to plated cell number and sorted live cells. **L.** Flow cytometry analysis of the colonies from K. **M.** Sort gating strategy. **N.** qPCR analysis of sorted D23 A1ATD1 MKs. Transcript levels for progenitor-associated transcripts were normalised to the *HMBS* levels and the levels in the MICB^-^PVR^-^ population. All error bars show standard deviation and replicates are shown as open points. **p <0.01, * 0.05 < p < 0.01, p ≥ 0.05. For all flow cytometry, viable cells were selected as DAPI negative and thresholds set on a corresponding isotype control for each sample.

We examined the transcripts enriched in the putative progenitor cluster and selected those that were highly expressed and cell surface associated as criteria for putative MKP markers. Among the potential markers (Figure 2b) we selected MHC Class I related peptide B (MICB) and Poliovirus Receptor (PVR). Details of their functions are included in Table 2. To investigate protein expression during differentiation, we carried out flow cytometry for cell surface MICB and PVR during MK differentiation (Figure 2c). Both MICB and PVR were expressed by higher proportions of cells on D9 of differentiation and decreased as the MKs reached maturity on D20 (as demonstrated by CD41 and CD42 expression). To gain better resolution on expression levels before D9, transcript levels of the markers were assessed by qPCR during early differentiation. *MICB* and *PVR* transcript levels peaked on D2 of differentiation and were reduced after this (Figure 2d) again indicating that they could be indicative of an early progenitor state.

**Table 1:**
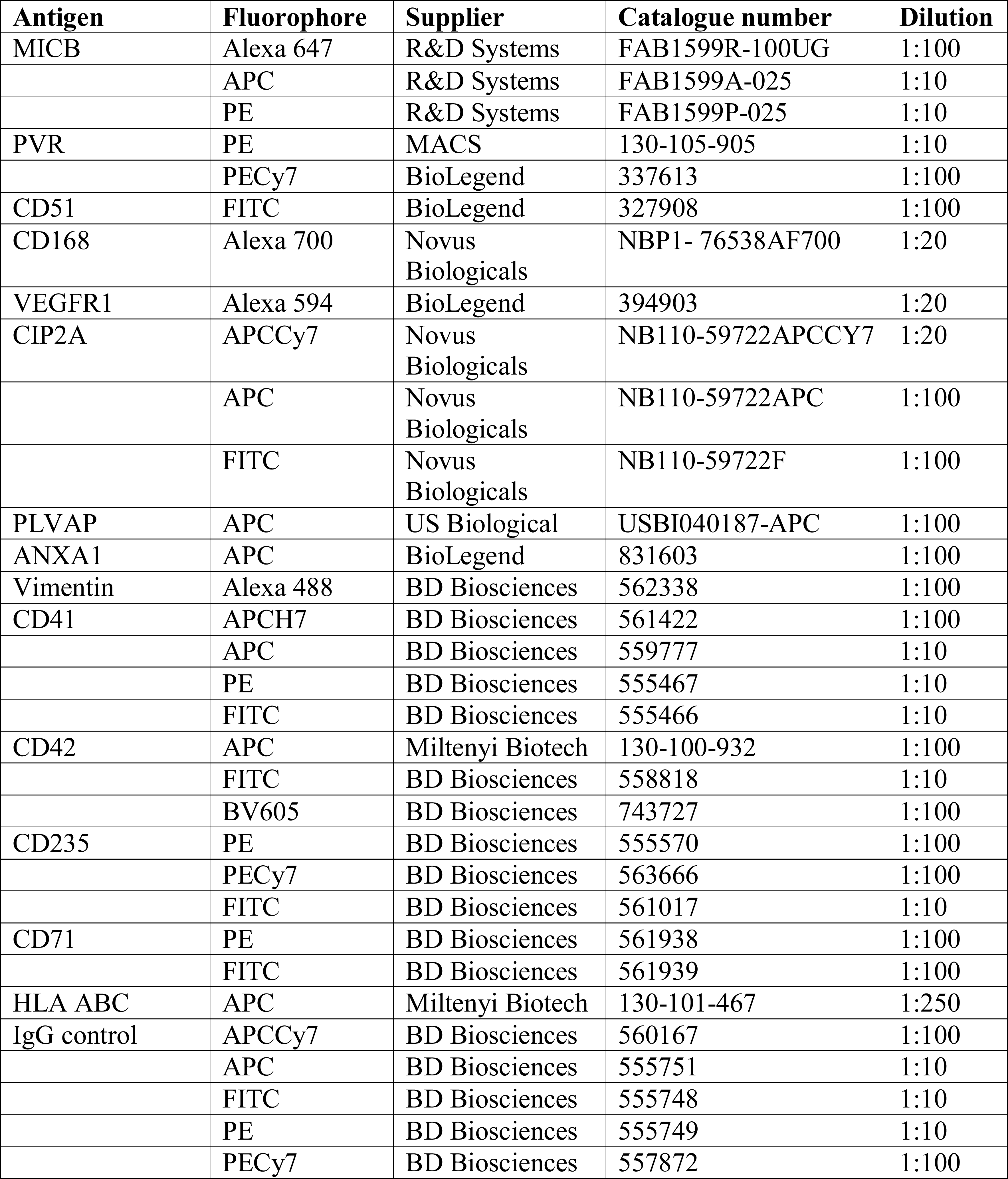
Antibodies.

**Table 2:**
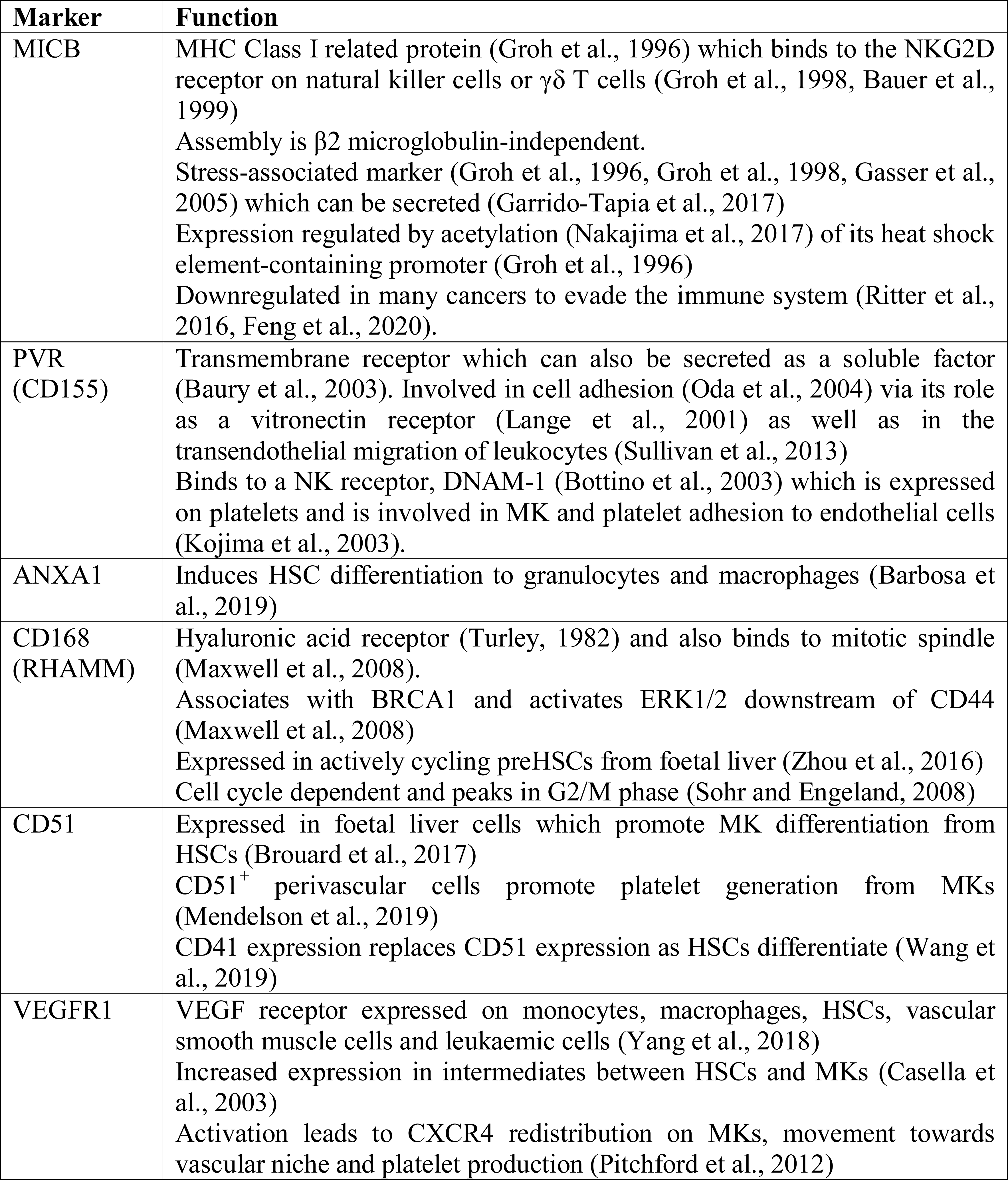
Putative progenitor markers.

To investigate whether these markers functionally define MKPs, we performed 20 index sorts (Figure 2e, STAR methods). MICB and PVR both strongly enriched for cells which formed colonies with p <2.2x 10^-16^ (Figure 2f). CD41 also strongly enriched for colony-forming cells (p <2.2x 10^-16^) whereas CD42 had a much weaker effect (p=0.08). The results of these index sorts indicate that cell surface expression of MICB and PVR is associated with MK colony-formation ability, and thus these two surface proteins could mark MKPs in the bulk culture.

To confirm this result, a Colony Forming Unit (CFU) assay was carried out (Figure 1d). MICB^+^ or PVR^lo^,^med^ or ^hi^ cells were sorted from a D75 A1ATD1 MK culture which was >86% CD41^+^ (Figure 2g). The sorted cells were plated in methylcellulose CFU assays and compared to live cells sorted without selecting for particular markers. Ten days later, colonies were scored (Figure 2h). MICB^+^ cells formed more than 4 times more colonies than live sorted cells, indicating that MICB marked MKPs. Interestingly, the level of PVR expressed also correlated with colony formation. PVR^hi^ cells formed more than twice the number of colonies in the control (Figure 2h, i). All colonies had consistent levels of CD41 expression when collected for flow cytometry analysis (Figure 2j), indicating robust formation of MKs. Colonies derived from MICB^+^ or PVR^hi^ cells expressed higher CD42 than other colonies (Figure 2j), indicating better maturation.

To ascertain whether sorting for both markers together would purify the progenitor population further, or whether both markers redundantly specified the same group of MKPs, cells were sorted for MICB and PVR in combination (Figure 2k). MICB^+^PVR^-^ cells formed fewer colonies than live cells, implying that all MICB^+^ colony forming potential is contained within the MICB^+^PVR^+^ fraction. Sorting for PVR enriched MKPs 2-fold and adding MICB to PVR further enriched colony formation to 3-fold. Flow cytometry analysis of the colonies (Figure 2l) demonstrated a similar mature MK production between conditions. We can conclude that adding MICB to PVR results in more efficient MKP purification from bulk MK cultures.

We then set out to measure the correlation of these sorted populations with the MKP state defined by the single cell sequencing. A set of transcripts were selected from the progenitor cluster shown in Figure 2a and qPCR primers were designed. D23 A1ATD1 MKs were sorted for MICB and PVR expression (Figure 2m) and progenitor-associated transcripts were quantified in these sorted populations by qPCR. Cells expressing either MICB or PVR alone expressed higher levels of progenitor-associated transcripts than control cells and MICB^+^PVR^+^ cells expressed even higher levels of progenitor-associated transcripts than either single positive population (Figure 2n). This further confirmed that using MICB and PVR together purified more progenitors than each marker used separately.

This first round of selection for MKP markers therefore allowed us to enrich MKPs by a factor of up to 4 (Figure 2k) using the CD41^+^MICB^+^PVR^hi^ gating strategy.

### Further refining MK progenitor markers: Single cell snapshot of established culture

*MICB* and *PVR* were very poorly detected on the time-course UMAP (Figure S4c and S4d). However, more highly expressed markers from this dataset could be used to further refine the MKP panel. Working from the time-course UMAP and focusing on the region we hypothesised to contain the MKPs, we selected the following markers: *ANXA1*, *HMMR* (CD168), *ITGAV* (CD51) and *FLT1* (VEGFR1) (Figure S4e-h). A summary of the literature surrounding these candidates is provided in Table 2. The markers were all more highly expressed in Clusters 6, 15 and 34 than the mature MKs (Clusters 5, 11, 48) on the time-course UMAP (Figure S4, Figure 1g, Figure 1j). These markers and combinations thereof should allow us to pinpoint the location of the MKPs: Clusters 6 and 34 are marked by *HMMR* and *ITGAV*, Cluster 15 by *ITGAV* and *FLT1* and Cluster 11 by *ANXA1* (Figure S4, Figure 1g). Additionally, the cluster projecting from cluster 6 has the highest homology to *in vivo* human MKPs and *HMMR* and *ITGAV* appear to be enriched in this area.

To determine whether these new markers were indeed expressed in MKPs, we purified CD41^+^ cells from a mature D27 A1ATD1 MK culture and sorted MICB^+^PVR^hi^ cells for sequencing using single cell 10x Genomics workflow with transgene expression quantification.

The 1661 cells which passed quality control were partitioned into 13 clusters and plotted on a UMAP (Figure 3a). We identified clusters of interest using the same approach as before and identified putative MKPs in Cluster 2 (enriched for cell division-associated GO terms and cell replication-associated transcripts such as *MCM3*, *5* and *7*, *PCNA* and *CENPW*), Cluster 1 (increased transcript levels of *KIT*) and Cluster 7 (high expression of *CD44* and *PLVAP*, both of which were enriched in the progenitor cluster of the Smart-seq scRNA sequencing). *MICB* and *PVR* were expressed by isolated cells from most clusters (Figure 3b, c), in keeping with the fact the cells were selected for high expression of both these genes. Transgene sequencing showed higher expression of *GATA*, *TAL1* and *FLI1* in clusters 1, 2, 7 and 8 (Figure 3d-f, Figure S6f). *ANXA1* marked Cluster 1 and was also weakly expressed in Clusters 7 and 8 (Figure 3g) whereas *HMMR* marked Cluster 2 (Figure 3h). *ITGAV* was broadly expressed at low levels (Figure 3i) whereas very little *FLT1* expression was detected, apart from a few cells in Cluster 1 (Figure 3j). Thus, by using a combination of all these markers (ANXA1, CD168, CD51 and VEGFR1) in index sorting, we aimed to pinpoint which clusters of both the progenitor UMAP and the time-course UMAP contained the MKPs.

**Figure 3:**
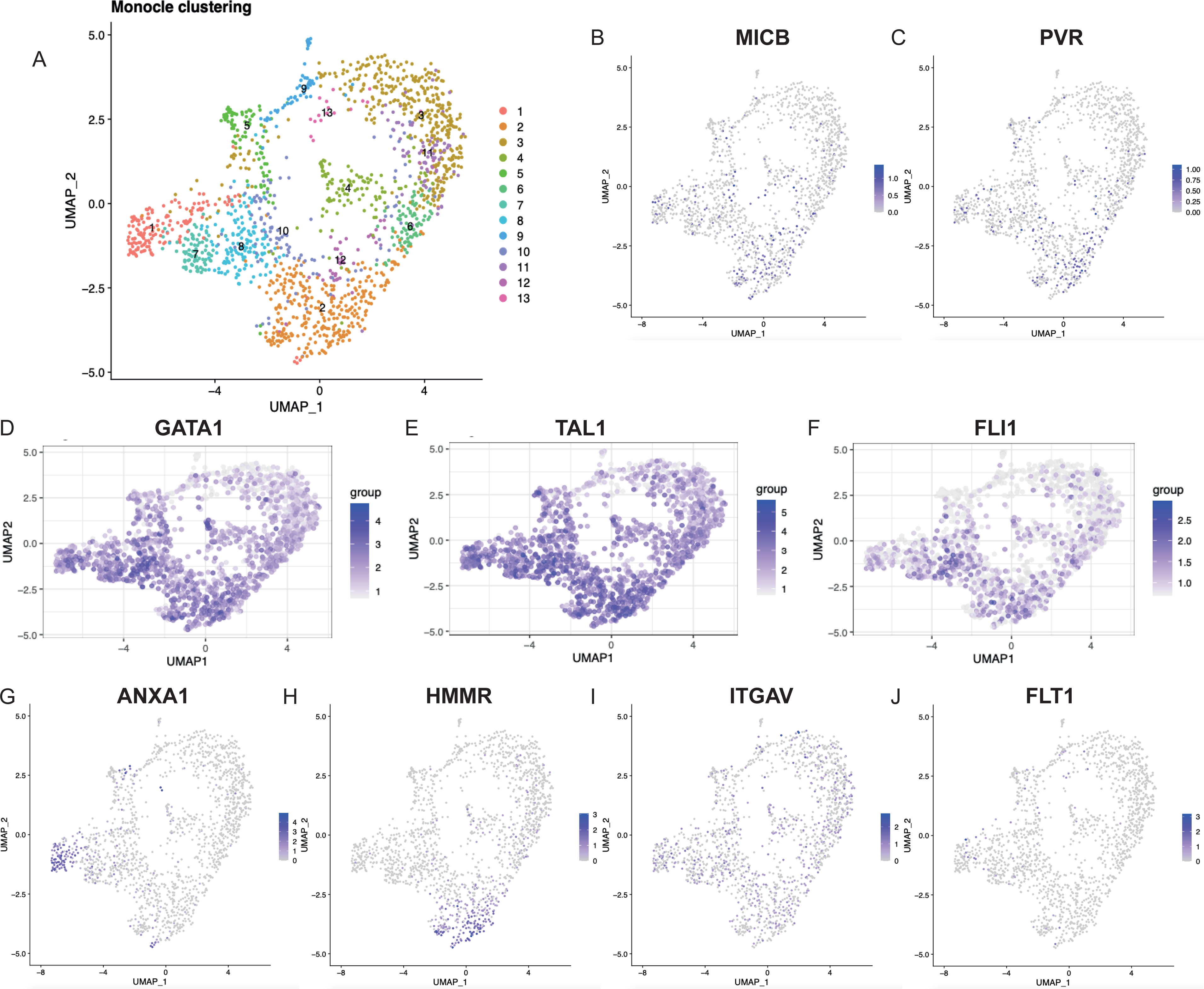
Single cell sequencing of megakaryocyte progenitors allows the selection of further candidate markers. **A.** UMAP of 10x Genomics single cell RNA sequencing of D27 A1ATD1 MKs sorted for top 50% MICB, top 26% PVR, bottom 33% CD42 expression. **B, C.** UMAP of single cell transform normalised *MICB* and *PVR* expression. **D-F.** UMAP of un-normalised *GATA1*, *TAL1* and *FLI1* expression from lentiviral transgenes plotted on the gene expression UMAP. **G-J.** UMAP of single cell transform normalised *ANXA1*, *HMMR*, *ITGAV* and *FLT1* expression.

### Random Forest feature selection identifies discriminative MKP markers

To investigate which combination of ANXA1, CD51, CD168 and VEGFR1 could select for MKPs and therefore identify the area of the UMAPs in which the MKPs were found, 20 index sorts were carried out on A1ATD1 MKs. The data was augmented to obtain a 1:1 ratio of positive and negative observations. Discriminative features were identified using the rank distributions from bagging analysis (Figure 4a). The most distinguishing marker for colony formation was PVR (median rank of 1 on 14 replicates) followed by MICB (median rank 1 on 4 replicates), confirming our previous colony formation results. CD41 and CD168 followed with median rank 1 on 1 replicate each. CD51 was median rank 2 on 2 replicates. (Figure 4a and Figure S5e). In contrast, ANXA1 performed poorly and did not appear to be associated with the MKP state. VEGFR1 was consistently ranked at the end of the selection table however it further purified populations selected by all other markers when used as a negative marker (p<0.05 between colonies and non-colonies by Wilcoxon test on the entire, normalised, outlier excluded dataset).

**Figure 4:**
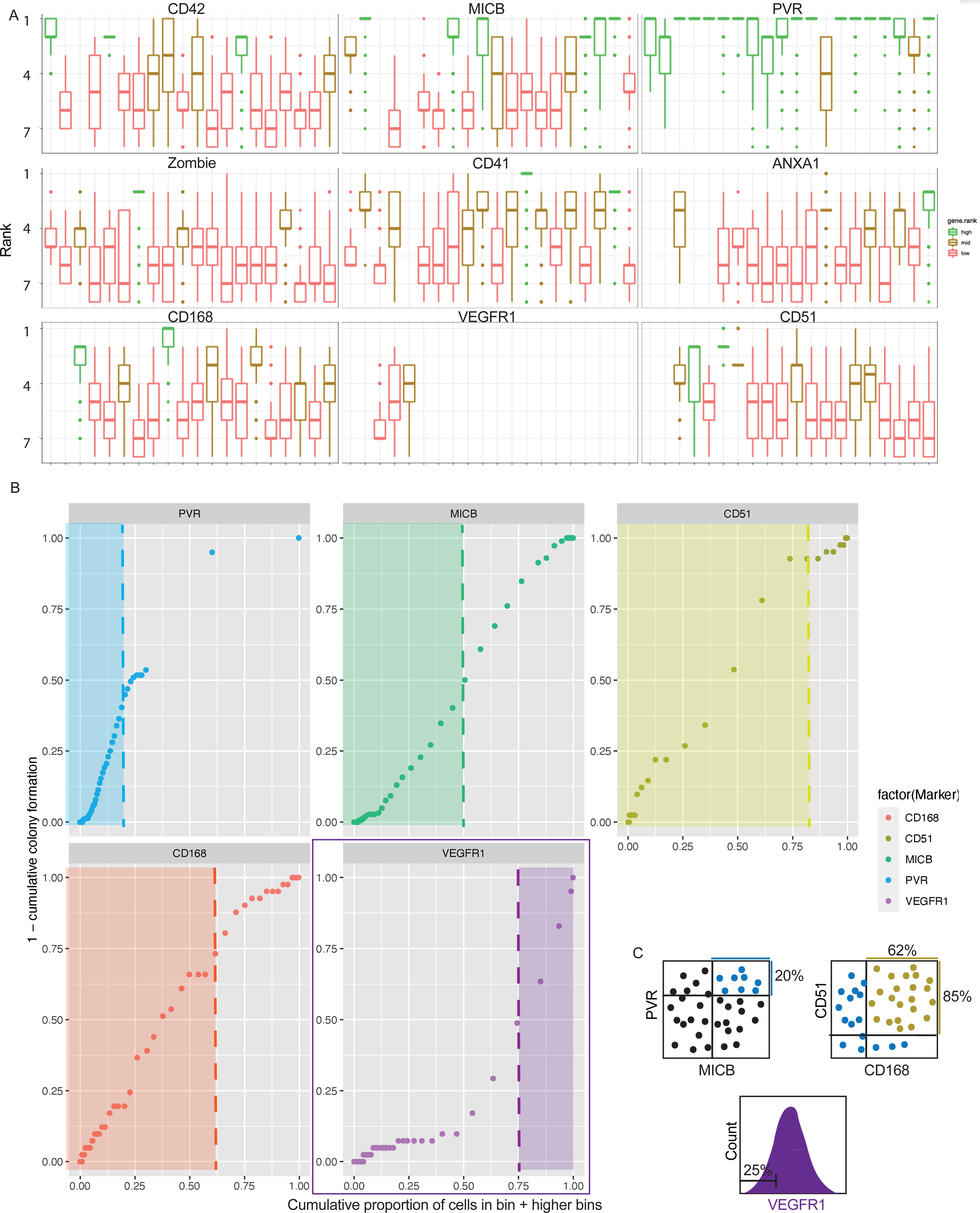
Identification of a panel of surface markers which can further purify MICB^+^PVR^+^ MKPs. **A.** 20 index sorts of A1ATD1 MKs. Rank indicates the first occurrence of a marker in a depth-first search of the decision trees making up the bagging model; genes with consistently low rank are discriminative for a given replicate. **B.** Determining cutoffs for MKP markers based on all index sorts of A1ATD1 MKs. Outliers were excluded and the 20 datasets were normalised and combined. The resulting dataset was then divided into 60 bins based on PVR expression and the colonies and non-colonies counted in each bin. This allowed the calculation of (1 - proportion colony formation per bin) (y-axis). Cells were then ordered with decreasing PVR expression and the proportion of cells with higher than this value of PVR plotted on the x-axis. The dataset was then subsetted on this expression level of PVR (top 20%) and the analysis repeated for MICB, CD51, CD168 and VEGFR1. Areas with maximum colony formation are shown with shaded boxes. **C.** Final sort strategy for MKPs. Cells are first purified as MICB^hi^PVR^hi^, then as CD51^hi^CD168^hi^ and finally as VEGFR1^lo^.

To determine the cutoffs required for most selectivity, all datasets were combined after outlier exclusion and normalisation. The combined dataset was binned into 60 bins and colonies and non-colonies scored in each bin, allowing the calculation of colony formation per bin. We then plotted the variable (1- cumulative colony formation percentage) as a function of (cumulative proportion of cells in each bin + bins with higher marker expression) (Figure 4b). At the bottom left of the graph are the cells with the highest expression level where 100% of cells form colonies. As we include cells with lower and lower levels of expression, the % of colony forming units decreases to the baseline corresponding to the bulk level, where all cells are included. The optimum cutoff of each marker is marked by the point of inflection of each graph. For positive markers we use cells below the cutoff but as VEGFR1 is a negative marker, we use the cells above the cutoff. As PVR was ranked the first marker in the highest number of index sorts, PVR was plotted first (Figure 4b). Selecting the top 20% of PVR expressing cells selected most MKPs. The dataset was then subsetted to contain only cells expressing this level of PVR and further refined using the other markers, in increasing order of p value from a single index sort. Selecting the top 50% of MICB-expressing cells, the top 85% of CD51-expressing cells, the top 62% of CD168-expressing cells and the bottom 25% of VEGFR1-expressing cells purified the highest proportion of total colonies. The final panel (Figure 4c) was subsequently tested on all datasets separately, outperforming all other marker panels generated by decision tree or random forest analysis.

Using the gene expression data of *ITGAV*, *HMMR* and *FLT1* (Figure S4), we then pinpointed the MKPs on the time-course UMAP (Figure 5a). These cells were located in clusters which lead to the mature MK cluster in trajectory analysis (Figure 1j), expressed highest *FLI1* levels among their timepoint (Figure 1k) and included the region of cluster 6 which was most similar to *in vivo* MKPs (Figure 1i). A similar analysis was carried out on the MICB^+^PVR^+^ UMAP (Figure 3) showing that the MKPs identified using this improved marker panel were found in cluster 2, which had most proliferation-associated GO terms (Figure 5b). Determining the surface marker expression of the MKPs allowed us to confirm their location on both UMAPs, pinpoint their emergence during differentiation and gain an insight into which transcripts define the MKP state.

**Figure 5:**
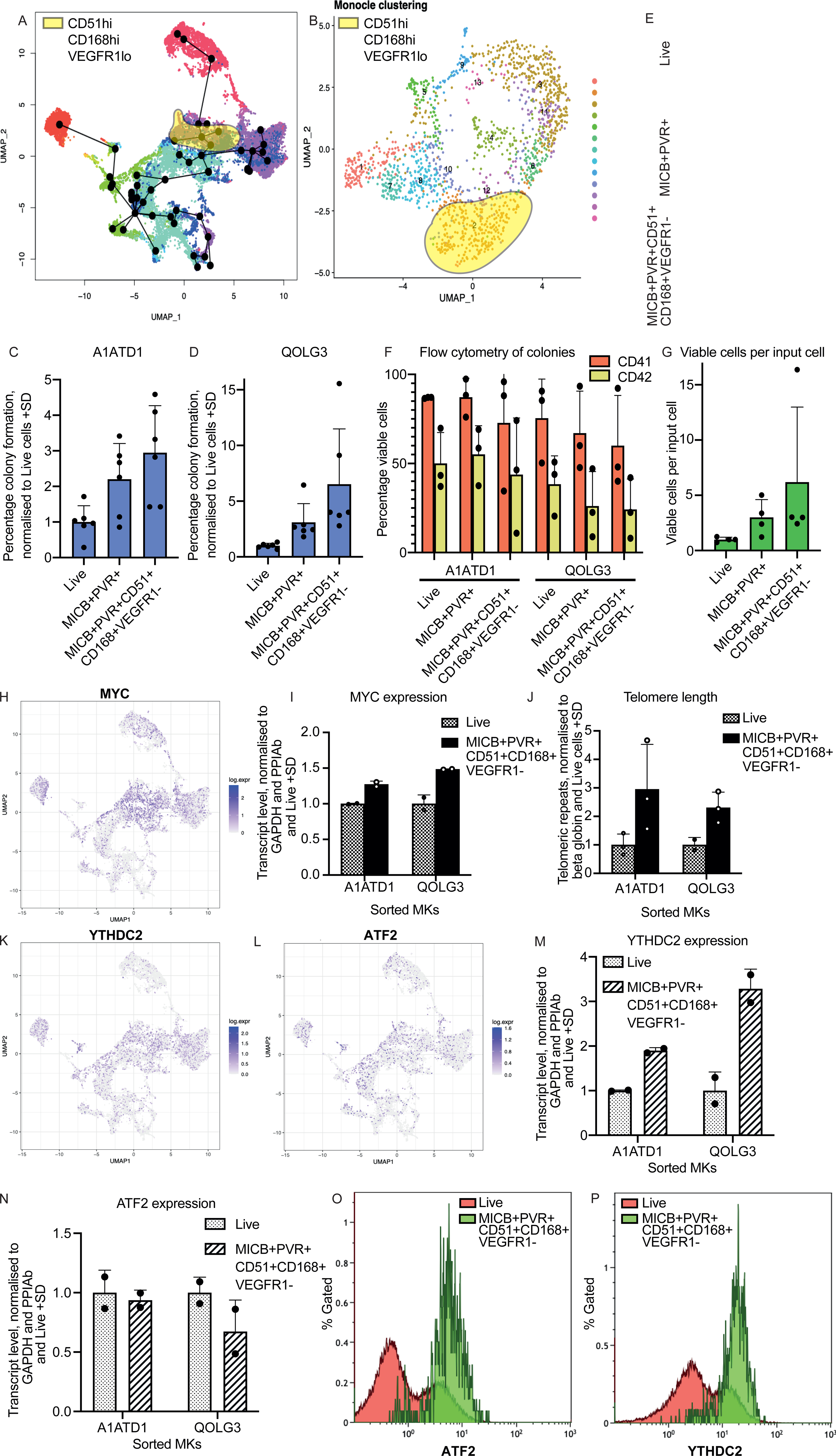
CD51, CD168 and VEGFR1 can further select MICB^+^PVR^+^ progenitors. **A.** *ITGAV*^hi^*HMMR*^hi^*FLT1*^lo^ cells’ location on monocle trajectory plot of differentiation. **B.** *ITGAV*^hi^*HMMR*^hi^*FLT1*^lo^ cells’ location on MKP UMAP. **C.** Colony counts from MK culture sorted as MICB^hi^PVR^hi^ alone (top 50% and 20% respectively) or in combination with CD51^hi^CD168^hi^VEGFR1^lo^ (top 85%, top 62%, and bottom 25% respectively). A1ATD1 MKs were sorted on D21, D27 or D33 in technical duplicate for each biological triplicate. Sorted cells were seeded in methylcellulose CFU assays. Colonies were counted 10, 9 or 11 days later respectively and normalised to total sorted cell number and the colony counts in the Live condition. **D.** Colony counts from MKs culture sorted as MICB^hi^PVR^hi^ alone (top 50% and 20% respectively) or in combination with CD51^hi^CD168^hi^VEGFR1^lo^ (top 85%, top 62%, and bottom 25% respectively). QOLG3 MKs were sorted on D24, D25 or D34 in technical duplicate for each biological triplicate. Sorted cells were seeded in methylcellulose CFU assays. Colonies were counted 11, 8 or 10 days later respectively and normalised to total sorted cell number and the colony counts in the Live condition. **E.** Brightfield microscopy images of colonies from C. Scale bar 500μm. **F.** Flow cytometry analysis of cells derived from the colonies in C and D. **G.** Total viable cell count after 2 A1ATD1 and 1 QOLG3 CFU assays. The entire well of colonies was collected and total DAPI negative cells per sample quantified using flow count fluorospheres. **H.** *MYC* expression levels on timecourse UMAP. **I.** qPCR analysis of transcript levels of *MYC* in a D27 A1ATD1 MK culture and a D34 QOLG1 MK culture sorted for live cells and for MICB^hi^PVR^hi^CD51^hi^CD168^hi^VEGFR1^lo^ cells. **J.** qPCR analysis of the quantity of telomeric repeats, normalised to globin in D21 A1ATD1 MKs or D34 QOLG3 MKs sorted as either live or MICB^hi^PVR^hi^CD51^hi^CD168^hi^VEGFR1^lo^.**K, L.** *YTHDC2* and *ATF2* expression levels on timecourse UMAP. **M, N.** qPCR analysis of transcript levels of *YTHDC2* **(M)** and *ATF2* **(N)** in a D27 A1ATD1 and a D34 QOLG1 MK culture sorted for live cells and for MICB^hi^PVR^hi^CD51^hi^CD168^hi^VEGFR1^lo^ cells. **O. and P.** Flow cytometry analysis of D38 QOLG1 MKs. Intracellular ATF2 **(O)**, YTHDC2 **(P)** and extracellular (MICB, PVR, CD51, CD168, VEGFR1) staining was carried out. Expression levels of ATF2 and YTHDC2 were plotted for total cells and MICB^hi^PVR^hi^CD51^hi^CD168^hi^VEGFR1^lo^ cells. For all flow cytometry, viable cells were selected as DAPI negative and thresholds set on a corresponding isotype control for each sample. For all qPCR, transcript levels were normalised to the geometric mean of *GAPDH* and *PPIAb* and the transcript level in the live sorted cells for two technical replicates of each biological replicate.

### Novel panel of surface markers reproducibly purifies MKPs from mixed cultures

We first tested whether this new panel (MICB^hi^PVR^hi^CD168^hi^CD51^hi^VEGFR1^lo^) outperformed MICB and PVR alone for MKP selection. MKs from two independent lines of iPSCs (A1ATD1 and QOLG3) were sorted. MICB^+^PVR^+^ cells showed increased colony formation in comparison to live sorted cells and the addition of the new markers further increased colony formation in both lines (Figure 5c, d, e). MK markers were similarly expressed in all conditions by flow cytometry of collected colonies (Figure 5f). Not only were there more colonies in the wells sorted for the new marker panel but these colonies were also larger than MICB^+^PVR^+^ derived colonies as evidenced by the much larger viable cell count in these wells as by flow cytometry (Figure 5g). Thus, combining CD51, CD168 and VEGFR1 with PVR and MICB can enrich MKPs by a factor of up to 6 fold in multiple iPSC lines. MKs derived from these progenitors have a surface marker profile which is identical to colonies from live sorted MKs, demonstrating that these progenitors generate mature MKs.

### MK progenitors have specific biology compared to mature MKs

The transcription factor MYC has been reported to act as a key rheostat for the MKP state (Feng et al., 2014). In iPSC directed differentiation protocols, MYC upregulation leads to the efficient generation of MKPs and its downregulation is required for platelet production (Takayama et al., 2010). MYC overexpression also generates immortalised MKPs which are capable of producing mature platelet-producing MKs upon MYC downregulation (Nakamura et al., 2014). We noted that *MYC* is upregulated in the clusters leading towards the mature MKs on the time-course UMAP and downregulated in mature MKs (Figure 5h). We confirmed this by sorting the population of MKPs using the markers described above and quantifying *MYC* expression by qPCR in two independent iPSC lines (A1ATD1 and QOLG3). MKPs expressed higher levels of *MYC* than live sorted cells (Figure 5i), consistent with their more proliferative state.

Telomeres are protective repeat sequences found at the end of chromosomes. Telomeres shorten as cells divide, which is well documented in haematopoietic cells (Vaziri et al., 1994). Patients with shortened telomeres can suffer bone marrow failure (Gramatges and Bertuch, 2013). As MKPs maintain the forward programmed culture for up to 3 months, we hypothesised that they should have longer telomeres than mature MKs. To investigate this, we sorted A1ATD1 and QOLG3 MKs for live cells and MKPs and carried out qPCR on the resulting gDNA for telomeric repeats, comparing them to the beta globin locus, of which there should just be two copies per cell. In both cell lines, MKPs had longer telomeric repeats than live sorted cells from the same culture (Figure 5j), indicating that these MKPs may have lengthened telomeres, supporting their long-term proliferative potential.

As MKPs exhibit a different transcriptional signature to mature MKs, we hypothesised that they may also have a different epigenetic profile and that our single cell data could provide insights into the mechanisms underpinning the MKP state. We therefore examined whether the MKP cluster identified in the time-course showed differential expression of epigenetic regulators. We found that *YTHDC2* and *ATF2* were both more highly expressed in the MKP cluster (Figure 5k and 5l). YTHDC2 is an RNA helicase which promotes the translation of m^6^A-modified mRNA (Mao et al., 2019) by attaching it to the ribosome (Kretschmer et al., 2018) and is essential for meiosis (Hsu et al., 2017). ATF2 is a histone acetyltransferase specific for H2B and H4 (Kawasaki et al., 2000) and regulates γ globin expression by histone deacetylase recruitment (Liu et al., 2013). Increased levels of both these epigenetic modifiers in MKPs could imply differential regulation of the translation of m^6^A-modified mRNA as well as increased histone acetylation at gene loci of key progenitor-related targets. We analysed the expression of *YTHDC2* and *ATF2* in the live cells and sorted MKPs from A1ATD1 and QOLG1 MK cultures and confirmed *YTHDC2* transcript levels were increased in MKPs, whereas *ATF2* transcript levels were unchanged (Figure 5m, n). However, when we analysed the expression of both these proteins by intracellular flow cytometry, both were increased in MKPs versus bulk MKs (Figure 5o and 5p). Therefore MKPs express higher levels of both these epigenetic modifiers than bulk MKs.

### Optimisation of culture condition for MKP production *in vitro*

Identifying specific markers for MKPs provided us with the tools to optimise culture conditions in order to promote MKP creation in early culture stages and the maintenance, proliferation and maturation of MKPs during long term culture. We first addressed the early stage of culture, carrying out MK differentiation under two oxygen (O_2_) concentrations; ambient (21%) and low (5%) O_2._ Low O_2_ has been shown to be beneficial for the culture of bone marrow-derived MKPs (Katahira and Mizoguchi, 1987). During MK differentiation from HSCs, low O_2_ was beneficial at early stages whereas ambient O_2_ was optimal for later stages (Lasky and Sullenbarger, 2011). We used three different concentrations of each of the cytokines SCF and TPO. The percentage of cells expressing CD41, CD42 and progenitor-associated markers were monitored along the differentiation time-course in this matrix of conditions.

More viable cells (Figure 6a) and more progenitors (Figure 6b) were produced on D9 in ambient O_2_ concentrations and these conditions retained the highest viable cell number (Figure 6c) and the most progenitors (Figure 6d) on D15. Low O_2_ conditions caused most MK maturation (Figure 6e).

**Figure 6:**
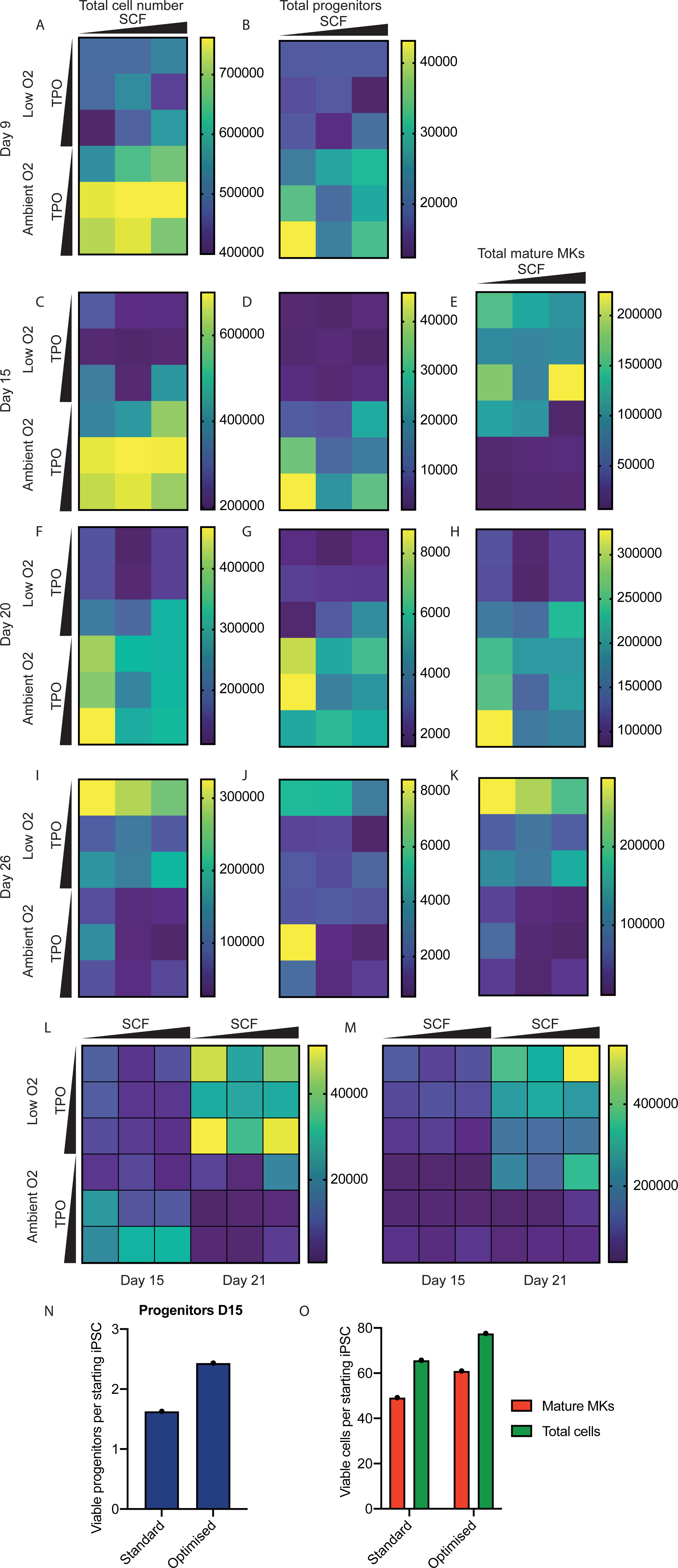
Optimising culture conditions for progenitor production *in vitro*. **A.** MK differentiation of QOLG3 iPSCs was carried out in varying concentrations of SCF (12.5, 25 and 50 μg/mL), TPO (10, 20 and 40 μg/mL) and O_2_ (5% (low) and 21% (ambient)). Total viable cell count per well (as defined by FSC/SSC and DAPI negativity, quantified by using flow count beads and flow cytometry). **B.** Total MICB^hi^PVR^hi^CD51^hi^CD168^hi^VEGFR1^lo^ cells (“progenitors”) were quantified in each condition. **C.** as A D15. **D.** as B D15. **E.** Total CD41^+^CD42^+^ cells (“mature MKs”) per well. **F-K.** Cell number, progenitor number and mature MK number on D20-D26. **L.** Total progenitor count in A1ATD1 MKs on D15 and D21. **M.** Total mature MKs per well in A1ATD1 on D15 and D21. **N. and O.** QOLG3 iPSCs were differentiated under standard and optimised cytokine and O_2_ concentrations. Number of progenitors **(N)** mature MKs and total cells **(O)** generated per starting iPSC in each condition. All samples gated on 21% O_2_, 25 μg/mL SCF, 20 μg/mL TPO.

On D20, the condition that had yielded most MKPs at D9 and D15 (ambient O_2_ and high TPO, low SCF) now yielded most MKs (Figure 6h), showing that early generation of MKPs results in the later generation of mature MKs. Ambient O_2_ still had most MKPs at D20 (Figure 6g), however low O_2_ promoted more MKP generation after D20 (Figure S6G). This increase production of MKPs in low O_2_ and low TPO after D15 translated into the highest number of mature MKs by D26 (Figure 6k), again demonstrating that increasing MKP generation early in the culture increased mature MK yield later on.

The experiment was repeated with A1ATD1 MKs and again more MKPs were produced on D15 at ambient O_2_, low SCF and high TPO but later on in culture, more MKPs were produced in low O_2_ (Figure 6l).

We then carried out an “optimised culture” where cells were kept at ambient O_2_, low SCF and high TPO up until D15 and thereafter in low O_2_ and low TPO. By D15, the optimised culture conditions improved MKP production to 1.5-fold (Figure 6n) and mature MK production 1.23-fold production over standard conditions (Figure 6o). In conclusion, the panel of surface markers identified as part of this study allowed us to quantify MKPs in culture and thus optimise cytokine and oxygen concentrations to maximise MKP production, thereby postponing culture exhaustion, maximising MK output and driving down the cost of platelet production.

## Discussion

In this study, we have mapped MK differentiation at high resolution from iPSCs to mature MKs, generating an overview of the cell fates acquired during differentiation and demonstrating that this protocol accurately generates MKPs and erythroid progenitors which mirror their *in vivo* counterparts in human bone marrow, peripheral blood and spleen. Microarrays have been used to analyse immortalised megakaryocyte lines (Nakamura et al., 2014, Ito et al., 2018) and RNASeq has been used to analyse initial MK differentiation from ESCs (Elcheva et al., 2014) and to compare MKs differentiated *in vitro* to MKs differentiated from cord blood-derived CD34^+^ HSCs (Moreau et al., 2016). However, this is the first time that *in vitro* produced MKs have been directly compared to *in vivo* progenitors. This study demonstrates that although *in vitro* MK differentiation appears to mirror *in vivo* development from pluripotent cells to mesoderm, haemogenic endothelium and finally commitment to the megakaryocytic lineage (Zhou et al., 2016), the cells do not pass through a state resembling HSCs or MPPs *in vivo*. Directed differentiation protocols often shepherd cells through a cell state characterised by the expression of CD31, associated with haemogenic endothelium (Feng et al., 2014, Mills et al., 2014) CD34, which marks HSCs (Takayama et al., 2008, Feng et al., 2014, Eicke et al., 2018). This study indicates that transcription factor-mediated forward programming may accelerate differentiation, proceeding directly to more committed progenitors.

Single cell sequencing has enabled us to pinpoint the transcriptional epi-states through which cells must transit to become mature MKs, using single cell trajectory analysis to identify clusters which correspond to MKPs and persist into late cultures. Confirming the results of previous studies (Dalby et al., 2018) and consistent with crucial role of FLI1 during megakaryopoiesis *in vivo* (Chen et al., 2014) and MKP differentiation from HSPCs *in vitro* (Zhu et al., 2018), we observe increased *FLI1* expression in the clusters leading to mature MKs by trajectory analysis. Combining index sorting for key marker candidates and machine learning on the resulting high-dimensional datasets, we have identified a panel of novel surface markers which can be used to purify MK progenitor cells as they differentiate from human iPSCs. Previous studies described panels to purify MKPs from MEP cultures (Psaila et al., 2016) or mouse bone marrow (Nakorn et al., 2003). This study describes the first surface marker panel allowing human MKP purification from iPSC-derived cultures, their quantification and the optimisation of their production.

The set of markers described here allowed MKP purification, providing insights into key aspects of MKP biology. We observed an increase in *MYC* expression in MKPs, supporting the results of other groups which describe the downregulation of MYC during MK maturation (Lu et al., 2018, Guo et al., 2009, Thompson et al., 1996, Nakamura et al., 2014, Bluteau et al., 2013, Chanprasert et al., 2006). Increased levels of MYC also explain the increased replicative capacity of MKPs. In addition, we also observed an increase in telomere length in purified MKPs, which may underlie their increased proliferative potential. As haematopoietic cells divide, either during aging or cytokine-supplemented culture, their telomeres shorten (Vaziri et al., 1994). This shortening can result in bone marrow failure, such as that seen during Dyskeratosis Congenita, aplastic anaemia and Fanconi anaemia (Gramatges and Bertuch, 2013). MKPs may require lengthened telomeres to support their replication for the extended periods of *in vitro* culture observed here.

Purification of MKPs using the marker panel described in this study also allowed us an insight into the epigenetic modifiers expressed by MKPs. We have described an increase in ATF2, a histone acetyltransferase for H2B and H4 (Kawasaki et al., 2000) in MKPs in comparison to cells in the bulk culture. Histone deacetylase inhibitors have been previously described to increase human CD34^+^ HSPC differentiation to MKPs (Zhu et al., 2018) indicating increased histone acetylation in the MKP state. ATF2 has been shown to upregulate the expression of γ-globin in erythroid progenitors (Liu et al., 2013) implying its involvement in erythroid differentiation. As MKPs are derived from the erythroid cluster of our time-course dataset, this could further underline the association of ATF2 with the MKP state. YTHDC2 is an RNA helicase upregulated in several cancers (Tanabe et al., 2014) which binds m^6^A-modified RNA (Hsu et al., 2017) and attaches them to the ribosome, “fast-tracking’ their translation (Kretschmer et al., 2018). Fewer granulocyte/erythroid/macrophage/megakaryocyte progenitors were formed by m^6^A depleted HSPCs in humans (Vu et al., 2017) and mice (Lv et al., 2018) indicating that m^6^A may be important for their differentiation. Interestingly, *MYC* RNA is marked by m^6^A in leukaemic cells (Weng et al., 2018, Vu et al., 2017) and m^6^A reduction reduces MYC (Vu et al., 2017) by reducing the half-life of its RNA (Weng et al., 2018). Therefore m^6^A is crucial for MYC expression. The increased levels of m^6^A binder YTHDC2 observed here may be linked to the increased *MYC* levels we observe in MKPs; YTHDC2 may stabilise *MYC* transcripts and expedition their translation.

Optimising cell culture conditions in order to increase cell output and diminish the cost-of-goods remains one of the major challenges in translating iPSC-derived cellular products to the clinic (Evans et al., 2021, Lawrence et al., 2021). Using the set of MKP markers identified here we found that adjusting oxygen and cytokine conditions can increase MKP generation, maximising mature MK generation. Ambient oxygen has been demonstrated to promote the later stages of MK differentiation from HSCs (Lasky and Sullenbarger, 2011). Interestingly, only later haematopoietic intermediates are generated in our differentiation system, explaining why ambient oxygen promotes optimal initial differentiation in our system. More progenitors were produced in later cultures under low oxygen tension in this study. Low oxygen tension has been shown to allow the culture of MKPs from human bone marrow (Katahira and Mizoguchi, 1987) implying that low oxygen can retain the MKPs already present in the cultures at this time. The optimised conditions identified here could thus allow the early expansion of MKPs and their retention over time.

In conclusion, the bioinformatic and biological tools illustrated in this study are broadly applicable to other stem-cell differentiation systems. They allow the identification of specific surface markers for cellular products, promoting a better biological understanding of stem cell types generated *in vitro*, allowing their comparison to *in vivo* counterparts and the optimisation of their production for the progress of novel stem cell-derived therapeutics towards the clinic.

## Limitations of study

This study focused on understanding the dynamics of MK differentiation from pluripotent stem cells. However, we do not know whether the MKP surface marker panel identified here can be applied to other MK differentiation systems such as directed differentiation or forward programming using alternative transcription factors. Similarly although the MKP transcriptome maps to human *in vivo* MKPs, it remains to be demonstrated whether the biological insights gained into MKPs’ epigenetic state and lengthened telomeres *in vitro* also apply to MKPs isolated from human bone marrow.

## Acknowledgements

M.L.’s research was funded by the UK Regenerative Medicine Platform, Pluripotent Stem Cell and Engineered Cell Hub (MR/R015724/1). This research was funded in part by a core grant from the Wellcome Trust (Grant Number: 203151/Z/16/Z) to the Cambridge Stem Cell Institute. For the purpose of Open Access, the author has applied a CC BY public copyright license to any Author Accepted Manuscript version arising from this submission.

The Wellcome 4-Year PhD Programme in Stem Cell Biology and Medicine funded the research of P.J.G (102160/B/13/Z), R.M. (222363/Z/21/Z), J.B. (MR/K500975/1) and M.S. (222398/Z/21/Z)

## Author contributions

M.L.: Conceptualization, Data curation, Formal Analysis, Investigation, Methodology, Validation, Visualization, Writing – original draft, Writing – review & editing. A.S.: Data curation, Formal Analysis, Methodology, Software, Visualization, Writing – original draft, Writing – review & editing. S.B.: Data curation, Formal Analysis, Methodology, Software, Visualization, Writing – review & editing. T.M.: Conceptualization, Data curation, Formal Analysis, Funding acquisition, Investigation, Methodology, Resources, Supervision, Validation, Visualization, Writing – review & editing. K.K.: Methodology, Resources, Writing – review & editing, M.P.: Methodology, Resources, Writing – review & editing. R.M., M.P., M.S., P.J.: Data curation, Formal Analysis, Investigation, Validation, Visualization, Writing – review & editing, J.B.: Conceptualization, Data curation, Formal Analysis, Investigation, Methodology, Software, Validation, Visualization, Writing – review & editing, C.P.: Conceptualization, Supervision, Writing – review & editing, I.M.: Conceptualization, Data curation, Formal Analysis, Investigation, Methodology, Project administration, Resources, Software, Supervision, Visualization, Writing – original draft, Writing – review & editing, C.G.: Conceptualization, Funding acquisition, Investigation, Project administration, Resources, Supervision, Writing – original draft, Writing – review & editing.

## Declaration of interests

The authors have no competing interests to disclose.

## STAR Methods

### Megakaryocyte forward programming

Induced pluripotent stem cells (iPSCs) [A1ATD1(Yusa et al., 2011), <u>HPSI1113i-bima</u>, <u>HPSI1113i-qolg_1</u>, <u>HPSI1113i-qolg_3</u>, <u>HPSI0813i-ffdk_1</u>] were seeded at single cell density on vitronectin (VTN-N, ThermoFisher A14700)-coated plates (Moreau et al., 2016). One day later, they were lentivirally transduced with *GATA1*, *TAL1* and *FLI1* in the presence of 10ng/mL protamine sulfate (Sigma, P4505) in DMEM/F12 (Gibco 11330-03), 0.05% NaHCO_3_ (Gibco 25080094), 65mg/L L-Ascorbic Acid (Sigma A8960) 1X ITS (Gibco 414000450) 10ng/mL BMP4 (Bio-Techne, 314-BP) and 20ng/mL FGF2 (Bio-Techne, 233-FB). The following day, cells were washed in PBS and the medium replaced without virus. On Day 3 (D3), the cells were washed in PBS and transferred to CellGenix GMP SCGM (CellGenix 20802-0500) supplemented with 25 ng/mL SCF (Gibco, PHC2116) and 20 ng/mL TPO (Bio-Techne, Cat.288-TP). On D10, the cells were removed from the plate using TrypLE Express (Gibco 12604021), centrifuged in PBS and replated in on uncoated tissue culture plates in CellGenix GMP SCGM supplemented with 25 ng/mL SCF and 20 ng/mL TPO. The culture was maintained at a density of 0.5-2 x 10^6^ cells/mL and cells were fed with SCF and TPO every 2 to 3 days by half medium exchange.

### 10x Genomics Single Cell sample preparation

Cells were dissociated at 24h intervals during the first 10 days of forward programming and washed twice in ice-cold PBS. D15 or D20 cells were counted and collected into ice-cold PBS, before being washed once more in ice-cold PBS. Cells were fixed in methanol, following the 10x Genomics demonstrated protocol for Single Cell RNA Sequencing (CG000136). They were rehydrated and submitted to CRUK CI Genomics Core Facility for library preparation using 10x Genomics Chromium Single Cell V(D)J Reagents Kits (protocol: CG000086 Rev H, chemistry 5’ v1.0: PN-1000014, PN-120236 and PN-1000020) with custom enrichment substituting the V(D)J primers for primers amplifying *GATA1*, *TAL1* and *FLI1* transgenes (Supplementary Table 1 and Figure S2a). Progenitor cells were stained and sorted as previously documented before being submitted for library preparation. A gene expression library and two enriched libraries (one for *GATA1*+*FLI1* and one for *TAL1*) were generated for each sample.

This experiment was sequenced in three batches; the library preparation was performed in two batches. The first run included samples D0_1, D1_1, D2_1, D3_1, D4_1, D5_1, D6_1, D7_1, D8_1, D9_1, D10_1,D15_1, and D20_1. The second run included all samples from the first run, except D8_1 and D9_1 which were replaced by D8_2 and D9_2. In addition, D5_2 was included as a bridging sample to allow the evaluation of batch effects. Two new samples, D0_single2 (D0_s2: iPSCs plated as single cells with ROCK inhibitor), and D0_timecourse2 (D0_t2: iPSCs plated with ROCKi) were also included. All sequencing data was generated using a NovaSeq6000. The first and second sequencing runs were performed with parameters Read1: 26bp, Index1: 8bp, Read2: 91bp. The third run included deeper sequencing of D3_1, D5_2, D8_2, D10_1, and D20_1 as well as all enriched libraries and was performed using PE150.

### Time-course 10x Genomics scRNA-seq analysis

Quality checks, alignment and protein-coding gene quantification were using Cell Ranger v3.1.0; to maintain consistency across sequencing runs in the time-course dataset, reads from the third sequencing run were aligned single-end with the R2 strands clipped to 91 bp; 72,310 cells were detected by Cell Ranger.

First the quality control was performed on the reads (Figure S2). The distribution of number of features per cell the sequencing depth per cell and number of cells per sample varied between samples. Consequently, the resolution of the time-course varied over time. The logarithmic relationship observed between sequencing depth and number of detected features per cell suggested that sequencing saturation was not reached.

Features (genes) expressed in fewer than 3 cells were excluded from downstream analysis; 23,949 genes were retained. Cells with <1,000 detected genes, >60,000 UMIs, or >10% reads incident to mitochondrial genes were discarded, leaving 22,236 cells for downstream analysis (Supplementary table 2). The dataset was normalised using sctransform.

UMAP (Uniform Manifold Approximation and Projection) of the filtered and normalised data (Figure S2) were used to assess the extent of batch effects between cells from different library batches and different sequencing runs. No clear separation was observed between cells from different sequencing runs, and while separation was observed between the library batches, the observed differences were likely the consequence of the overrepresentation of certain timepoints in either library batch. Based on these results, no batch correction was deemed necessary.

UMAPs coloured by raw and normalised sequencing depth were inspected to check whether the heterogeneity observed in the dimensionality reduction was confounded with sequencing effects. Raw sequencing depth was uneven across the UMAP, but normalisation reduced this variation across the cells. Normalised mitochondrial and ribosomal protein-coding expression levels were plotted on the UMAP and regions generally expressed relatively high levels of either but not both.

After normalisation using sctransform, clustering was performed with Seurat v3.1.4 (SLM algorithm (Waltman and van Eck, 2013) on the 5-nearest neighbor graph), Monocle v2.14 (density-peak based clustering on a t-SNE dimensionality reduction), and PCA followed by k-means, on the top x=(500, 1,000, 2,000) most abundant genes. Element-centric similarity (ECS, Gates et al., 2019) was used to assess the stability of the clustering with regard to the number of genes. The Monocle clustering on the 1,000 most abundant genes, which comprised between 60% and 90% of counts in cells across samples, displayed the highest stability; these genes were subsequently used for dimensionality reduction using PCA and UMAP (created on 30 PCs, selected after inspection of elbow plot). The stability of the clustering, in terms of number of clusters was assessed using the proportion of ambiguously clustered pairs (Oenbabaoğlu et al., 2014). Cells from each sample are often clustered together in a small number of clusters while individual clusters often extend beyond a single time point sample (Figure S2m). Marker genes for each cluster were identified using the ROC test in Seurat; genes with |ln(FC)| > 1 were considered for the ROC test; the selected-markers for each cluster were the top 25 genes ranked by discriminative power (Supplementary Table 3).

Gene set enrichment terms were calculated with g:profiler (Reimand et al., 2016), using the 25 genes with highest power in the ROC test, with all genes returned by sctransform across all clusters used as background (Supplementary Table 4). A blacklist of biologically unrelated annotation terms was applied to avoid inflated similarities. Enrichment terms with g:SCS-adjusted p-values <0.05 were further explored.

The cell cycle phase of cells was computationally inferred using the CellCycleScoring function in Seurat, using an annotated list of cell cycle-associated genes (Tirosh et al., 2016).

Pseudotime trajectories were calculated using Monocle on a DDRTree dimensionality reduction on the Pearson residuals of the abundant genes with parameter max_components=2, and separately using Slingshot v1.4 (Street et al., 2018) with default parameters on the original UMAP representation. Temporally expressed genes were identified with quadratic regression of each gene’s expression on the Slingshot pseudotime variable. The heatmap was created using the 100 most significant genes, where, for ranking purposes, the smallest of the two p-values for the linear term and the quadratic term was used for each gene.

### *In-vivo* 10x Genomics scRNA-seq integration

An *in-vivo* 10x Genomics scRNA-seq dataset of human haematopoietic stem and progenitor cells from the bone marrow, peripheral blood and spleen of two donors was used to assess similarities between *in-vivo* haematopoietic intermediates and *in-vitro* megakaryocyte differentiation using our optimised protocol. We used as starting point the count matrix with raw expression levels and metadata with annotated cell type identities as presented in Mende et al., 2020. Based on inspection of distributions of sequencing depth, number of detected features, proportion of UMIs assigned to mitochondria and ribosomal genes (MT% and RP%, respectively), cells with >1,000 features, <10% MT UMIs and >20% RP UMIs were retained for further analysis, totalling 13,099 cells for donor 1 and 17,350 cells for donor 2. After normalisation with sctransform we used the dimensionality reductions, PCA and UMAP (created on 30 first PCs based on elbow plot of variance per component), to assess the consistency between patients. Substantial batch effects were observed between the two donors; consequently normalisation and analysis of the data was performed separately per donor to preserve donor-specific characteristics. Subsequently, only cells annotated as megakaryocyte erythroid progenitor, haematopoietic stem cell/multipotent progenitor, megakaryocyte progenitor, or erythroid progenitor were used for downstream analysis, leaving 9,205 donor 1 cells and 12,214 donor 2 cells.

To identify marker genes for the annotated cell types, the ROC test in Seurat was used with minimum fold-change of 2.0; taking the union of positive markers across each cell type in both donors, 101 discriminative genes were detected. Random forest models (Breiman, 2001) were trained separately per donor using the expression of these 101 genes and the annotated cell types as output (using the randomForest R package, version 4.6-14). 100 decision trees were used in the RF models; the convergence of the models was assessed using training error versus number of trees; the models converged after 40 trees. The stability of the RF models was assessed using 10-fold cross-validation; for the RF trained on donor 1, the distribution of accuracies was high (min 92.7%, median 93.4%, max 95%, stddev 0.7%), and the RF trained on donor 2 it was similarly accurate (min 90.3%, median 91.3%, max 93.4%, stddev 0.9%).

Next, using the RF model, we predicted the cell types for the 10x Genomics scRNA-seq time-course. A cell is assigned to the cell type with largest probability; by applying a minimum probability threshold, uncertain cell type predictions can be avoided. To determine the probability threshold, the percentage of D0 cells assigned to any cell type was used as criterion; the lowest probability threshold that ensured <1% of D0 cells were assigned to any type (as D0 cells should be in the primed pluripotent state and not represented in adult human bone marrow, spleen or peripheral blood). For donor 1 the threshold was 0.53 and for donor 2 the threshold was 0.67.

### Sample preparation and Smart-Seq of mature megakaryocyte culture

Single cells from a Day 44 A1ATD1 MK culture were sorted into two 96-well plates containing 2.3μL 0.2% Triton X-100 with 1X RNase inhibitor and frozen to −80°C before library preparation. Reverse transcription was initiated with oligo-dT primers and SuperScript-II reverse transcriptase (ThermoFisher Scientific, 18064022) and supplemented with ERCC RNA Spike-In. The cDNA was then PCR amplified for 21 cycles using the KAPA HiFi Hotstart ReadyMix and IS primers (Kappa Biosystems, KK2601). After purification of the amplified cDNA, the indexed library was made using the Illumina Nextera XT DNA kit. The libraries from 96 cells were pooled and quality was checked on an Bioanalyser high sensitivity DNA chip (Agilent, 5067-4626) before sequencing on one lane of an HiSeq 4000 (Illumina).

Quality control of raw reads was performed using FastQC v0.11.3 and summarised with MultiQC v1.8. Checks included variability of sequencing depth and distribution of GC content. Samples were aligned to the GRCh38.p13 reference genome with STAR v2.7.3a (default parameters, (Dobin et al., 2012) using Ensembl v98 annotations. Protein-coding gene quantification was performed using featureCounts v2.0 (default parameters). Scatterplots of sequencing depth versus number of detected features per cell were visualised; cells with fewer than 200,000 reads were excluded, leaving 123 cells for downstream analysis.

Normalization of raw gene abundances was performed using sctransform (Hafemeister and Satija, 2019).

Across the 192 samples, the distribution of raw sequencing depth was wide (Figure S5); after normalisation and cell filtering, the distribution narrowed. We infer that sequencing saturation was reached for this experiment based on the lack of linear or logarithmic relationship between sequencing depth and number of features. We conclude that the observed heterogeneity in the data is not primarily dependent on the variation in sequencing depth.

To select the number of abundant genes, cluster stability with regard to gene expression was assessed using element-centric clustering. Using the density decision graph (Figure 2a inset), we identify only three cells that exhibit high density and are distant from cells of higher density; these cells are subsequently assigned as cluster centers (Rodriguez and Laio, 2014).

Subsequently the 2000 most abundant genes, accounting for between 60% and 80% of counts, in a majority of cells, were used for PCA, and UMAP (created on the 30 first PCs were used after inspection of elbow plot), as well as clustering. Two clustering approaches were compared side by side: graph modularity optimization-based clustering with Seurat v3.1.4 (Butler et al., 2018) and density peak-based clustering with Monocle 2.14 (Trapnell et al., 2014). A nearest-neighbour graph with shared nearest neighbour weighting of the edges was created with Seurat using the first 30 PCs, with k=20 neighbours; this was used for clustering using the SLM algorithm (Waltman and van Eck, 2013). Clustering was separately performed with Monocle v2.14, using a tSNE dimensionality reduction and the density peak algorithm (Rodriguez and Laio, 2014). Monocle was configured to find the same number of clusters (3) as Seurat (for the latter, detected with default resolution parameter). Marker genes for each cluster were identified using the ROC test in Seurat. Only genes with |ln(FC)|>1 were considered for the ROC test, and the final markers for each cluster were the top 20 positive markers ranked by discriminative power. Potential progenitor markers and the GO terms associated with cell surface expression were selected through manual curation.

### Sorted megakaryocyte progenitor 10x Genomics scRNA-seq

A D27 A1ATD1 MK culture was sorted for cells expressing PVR (top 26%) and MICB (top 33%). Live cells were subjected to 10x Genomics library preparation, generating separate libraries for gene expression and transgene expression as detailed previously. Pre-alignment quality control was performed analogously to the smart-seq2 analysis; subsequently the 10x Genomics GRCh38 v3 reference genome was used for alignment and protein-coding gene quantification with Cell Ranger v3.1.0; 1899 cells were detected. The number of features and counts varied across cells; MT and RP distributions prior to filtering displayed long tails (Figure S6). The logarithmic relationship between number of UMIs and number of features detected per cell suggested sequencing saturation was not achieved. After quality control and filtering, the distribution of counts per cell was narrower. UMAPs of MT and RP indicated that groups of cells have a high proportion of either MT or RP but not both.

Features expressed in fewer than 3 cells were excluded from further analysis; 16,660 genes were retained. Distributions of sequencing depth, number of detected features, as well as percentages of mitochondrial (MT) and ribosomal protein (RP) incident reads per cell were inspected; based on these distributions, cells with <1,000 unique features or >10% MT reads were discarded; 1661 cells were retained for downstream analysis.

Sctransform was used for normalisation of expression levels across cells; based on the element-centric clustering similarity across clustering methods, 2000 most abundant genes, accounting for between 70% and 90% of counts in a majority of cells, were used for dimensionality reductions (PCA and UMAP created on 30 first PCs) and clustering; for each iteration of the pipeline we chose the highest number of genes accounting for the target percentage of assigned counts. Seurat and Monocle were used for clustering with the same parameters as for the Smart-seq2 data (Supplementary Tables 6 and 7), with the adjustment that the kNN graph for Seurat used the 5 nearest neighbours. Imputation of expression levels was performed with ALRA (Linderman et al., 2018); the imputed values were only used for UMAP visualisation.

### Flow cytometry

Megakaryocytes were stained with the corresponding antibodies (Table 1) in PBS/0.5% BSA/5mM EDTA for 20min at room temperature. They were then washed in PBS/0.5% BSA/5mM EDTA, centrifuged for 8 min at 120g and resuspended in PBS/0.5% BSA/5mM EDTA with 1X DAPI. They were analysed using a Gallios flow cytometer (Beckman Coulter A94303) or sorted.

### qPCR

RNA was extracted using an RNeasy Mini Plus kit (Qiagen 74104) with gDNA Eliminator columns. cDNA was synthesised using a Maxima First Strand cDNA Synthesis Kit (Thermo Scientific K1641). qPCR was carried out using Applied Biosystems Fast or Power SYBR Master mix, 0.5μM primers (Supplementary Table 1) on an Applied Biosystems StepOne thermocycler. Genomic DNA for qPCR was extracted using a Wizard SV kit (Promega A2361) and analysed as above.

### Index sorting

Single cells were index sorted (Wilson et al., 2015) using a FACS Aria II (BD Biosciences) or a FACSAria Fusion (BD Biosciences) into flat bottomed non-coated 96- or 384-well plates which had been filled with 100μL/30μL CellGenix GMP SCGM supplemented with 25 ng/mL SCF, 20 ng/mL TPO and 1X Pen/Strep. The plates were incubated for 10-14 days at 37°C, 5% CO_2_, 21% O_2_. The colonies were then counted and imaged.

### Bioinformatics methods for processing the Flow cytometry output

Wells containing a colony of more than 3 megakaryocytes on D9-14 of culture were scored as positive observations; colonies containing fewer than 3 cells were scored as negative. To determine genes that were expressed differently in positive and negative observations, the Wilcoxon rank-sum test was applied. The distribution of gene values and scatterplots of intensities of each gene pair, were visualized, as well as PCAs of the data. Standard logistic regression and random forest models (the latter using 1,000 decision trees) were used to build classifiers for the observations; both sets of models failed to classify positive observations due to their low frequency (randomForest R package v4.6-14 (Breiman, 2001)).

Across replicate plates, we observed considerable variability in the number of colonies. Two replicates, containing 2 and 4 colonies respectively, were excluded from data augmentation and the bagging analysis due to their low number of colonies. Synthetic positive observations were generated using SMOTE (Chawla et al., 2002), to create an augmented dataset with 1:1 ratio between positive and negative observations (no down sampling of negative observations was applied). The distributions of gene expressions in the augmented and original data were visualised and compared to assess changes arising from augmentation. Using the augmented data as training set, both standard logistic regression and random forests succeeded in classifying positive observations on the original dataset, at the cost of a higher false positive rate for logistic regression.

To find the most discriminative features for separating positive and negative observations, the decision trees making up the random forest models were used. Specifically, the distribution of gene intensities used for splits and the ranks of a gene where it is first used in the tree were visualised. Genes with a consistently high rank (indicating they are used early in trees to separate observations) and a tight distribution of intensities (indicating that the gene consistently separates observations at the same expression level) were considered good candidates for improvement of the culture system. Separately, negative observations were subsampled to 30% of their original frequency to assess logistic regression, bagging decision trees, and random forest (the latter two using 1,000 decision trees and otherwise default parameters) performance when trained on subsampled data; false positive and false negative rates were consistently higher than the corresponding model trained on augmented data, and subsampling was not used further.

To identify the most discriminative features for separating positive and negative observations/ colonies, the decision trees making up the bagging models were used. Specifically, when traversing the trees depth-first, the rank of the first occurrence of genes was recorded; subsequently the distributions of ranks per gene and dataset were visualised using histograms and boxplots. Genes with consistently high rank across datasets were considered good candidates for improving the culture system. The rank distributions were summarised by calculating the means and medians of the ranks per gene and dataset, and the distributions of means and medians per gene were inspected.

### Methylcellulose Colony forming assay

Single cells were index sorted (Wilson et al., 2015) using a FACSAria II (BD Biosciences) or a FACSAria Fusion (BD Biosciences) into round bottomed BD Falcon tubes containing PBS/0.5% BSA/5mM EDTA. The tubes were centrifuged at 120g for 8min and cells were resuspended in 300uL basal medium and added to 3.3mL non-enriched MethoCult (Stemcell Technologies, H4535) supplemented with 50ng/mL Stem cell factor (Gibco, PHC2116), 100ng/mL thrombopoietin (R&D, 288-TP) and 1X PenStrep (Gibco, 15140122). 1.2mL was plated in duplicate wells of a SmartDish (Stemcell Technologies, 27370). Plates were incubated at 37°C, 5% CO_2_, 21% O_2_ and colonies were counted after 8-14 days using an Olympus brightfield microscope.

## Supplemental information titles and legends

**Figure S1: Sorting megakaryocytes for CD34, KIT, CD61 or KDR does not enrich for MKPs. A.** Flow cytometry plots of CD34, CD42, CD41, CD235 and KIT expression in a D45 A1ATD1 culture was sorted for CD34 and KIT levels. Sorted cells were seeded in a methylcellulose colony formation assay, supplemented with SCF and TPO to promote MK formation. The resulting colonies were counted (central graph) and analysed by flow cytometry (pie charts), selecting viable cells by DAPI negativity and gating on isotype controls for each antibody. **B.** Flow cytometry strategy for sorting D29 A1ATD1 MKs for CD61 or KDR expression. Colony count from MKs sorted for the markers shown and plated in a methylcellulose colony formation assay, supplemented with SCF and TPO. Colonies were scored on D15. **C.** Transgene expression strategy for 10x Genomics sequencing. V(D)J primers from the target enrichment library preparation protocol were replaced with primers for regions in the lentiviral backbone, allowing libraries to be generated of the overlap regions between transcript and lentiviral backbone, ultimately providing a readout of transgene expression levels. LTR: Long Terminal Repeat. EF1α: Elongation Factor 1α promoter, WPRE: woodchuck hepatitis virus posttranscriptional regulatory element

**Figure S2: Time-course quality control summaries suggest that biological variation and not technical effects are the primary driver of post-normalisation transcriptional heterogeneity. A-D.** Distributions of number of features (A), unique molecular identifiers (UMI) (B), mitochondrial (MT) reads % (C), and ribosomal protein (RP) reads % (D) across cells, split per sample illustrate differences across the time points. **E.** Scatter plot for evaluating sequencing saturation. The logarithmic relationship observed for UMI counts versus number of detected features per cell suggest that sequencing saturation was not reached. **F.** Histogram illustrating the number of captured cells per sample (target cells per time point: 2000). The observed variation underlines the heterogeneity in resolution across the time-course; note the y-axis is on a log10 scale. **G.** Batch summary across the time points. UMAP of cells coloured by biological batch illustrates some overlap across the two batches, suggesting that the dimensionality reduction is not driven primarily by biological batch. Inset panel: D5 samples across batches. **H.** Sequencing run summary across the time points. UMAP of cells coloured by sequencing run illustrates an acceptable overlap and suggests no significant batch effect. **I-J.** UMAPs coloured by raw (I) and normalized (J) sequencing depth illustrate that normalization reduces differences across UMAP regions. **K-L.** UMAPs presenting the MT% (K) and RP% (L) across cells using a colour gradient illustrate regions of UMAP space high in MT% or RP%; these regions are, in their majority, disjunct. **M.** UMAP faceted by cell origin, coloured by Monocle cluster. Samples corresponding to the same timepoint but different sequencing run or biological batch are similar both in location in UMAP space and in cluster membership. Example: D5.2: Day 5, batch 2.

**Figure S3: Identification of cell identity clusters on the UMAP embedding of the single cell RNA sequencing of megakaryocyte differentiation. A.** 51 Monocle clusters shown on the UMAP of the MK differentiation process. **B and C.** Normalised expression levels of transcripts associated with the naïve (B) and primed (C) pluripotent state in the monocle clusters shown in Figure S3A corresponding to iPSCs. **D.** sctransform-normalised *HAND1*, *KRT8*, *KRT18*, *KRT19*, *CD34*, *KDR, CDH5, PECAM*, *GATA1*, *TAL1*, *FLI1*, *GYPA, ITGA2B* and *GP1BA* expression shown in on the UMAP the MK differentiation time-course. **E.** Random forest-predicted cell types on single cell time-course data. Cell types learned on 10X haematopoietic stem and progenitor cells isolated directly from human bone marrow, spleen and peripheral blood^57^; model deployed here trained on donor 2: 3% D0 cutoff, threshold of 0.62. Non-overlapping groups of MEP and MkP are observed during *in vitro* MK differentiation. MEP: Megakaryocyte Erythroid Progenitor, HSC/MPP: Haematopoietic Stem Cell/Multipotent Progenitor. MkP: Megakaryocyte Progenitor, EryP: Erythroid Progenitor. H: Donor 2: 1% D0 cutoff, threshold of 0.667.

**Figure S4: Expression of putative megakaryocyte progenitor markers during megakaryocyte differentiation A.** Heatmap of most significant temporally changing genes defining the monocle trajectory shown in Figure 1j. Each row shows the expression of one gene on a random subset of cells across the pseudotime trajectory. **B.** Unnormalised *GATA1*, *TAL1* and *FL11* expression shown on the UMAP of the scRNA-seq experiment of the MK forward programming time-course in enriched libraries and gene expression library. **C-H.** Normalised *MICB*, *PVR*, *ANXA1*, *HMMR*, *ITGAV* and *FLT1* expression shown on the time-course UMAP (Figure 1f).

**Figure S5: Quality control summaries of Smart-seq data. A.** The distribution of unnormalized sequencing depths (ncount) show high variation across cells.; cells with ncount > 200,000 are further investigated. **B.** Distribution of normalised (using sctransform) sequencing depths per cell post-filtering; a tighter distribution of normalised ncounts is obtained; no further filtering is applied. **C-D.** Scatterplots of sequencing depth versus number of detected features per cell that indicate sequencing saturation; values on the x-axes are on linear (C) and log2 (D) scale; values on the y-axes are on linear scale. **E.** Boxplots of bagging analysis, performed on repeatedly subsampled data from 20 index sorted A1ATD1 MKs stained for the markers shown on the x axis.

**Figure S6: Megakaryocyte progenitor cell sequencing quality control summaries and the optimisation of MKP generation by cytokine and oxygen titration A.** Distributions of unique molecular identifiers (UMIs), number of detected features, % of reads incident to mitochondrial genes (MT) and ribosomal proteins (RP) illustrate variability among cells prior to filtering and normalization; cells with nfeature < 1,000 or MT% > 10% were discarded from downstream analysis. **B.** Scatter plot for evaluating sequencing saturation. The logarithmic relationship observed between the number of detected features versus UMI counts per cell suggests sequencing saturation has not been achieved. **C.** Distributions of UMI counts, number of features, MT% and RP% post filtering and normalisation. The tighter distributions suggest an increased comparability of the cells. **D-E.** UMAP embeddings presenting the MT% (D) and RP% (E) across cells using a colour gradient illustrate regions of UMAP space high in MT% or RP%; these regions are, in their majority, disjunct. **F.** Unnormalised *GATA1*, *TAL1* and *FL11* expression in enriched libraries and gene expression library shown on the UMAP embedding of the scRNA-seq of sorted MICB^+^PVR^+^ MKPs. Two enriched libraries were sequenced for each transgene combination. **G.** Change in cell number, progenitor number and mature MK number on D15-D26 of QOLG1 MK differentiation. Data shown in Figure 6 with each timepoint normalised to the previous timepoint.

**Supplementary Table 1: Sequences of primers used for qPCR and 10x Genomics enriched libraries amplification.**

**Supplementary Table 2: Samples used for single cell RNA sequencing of *in vitro* megakaryocyte differentiation time-course.** Statistics are shown pre- and post-filtering.

**Supplementary Table 3: Markers of each monocle cluster from time-course single cell RNA sequencing experiment.**

**Supplementary Table 4: GSEA terms associated with each monocle cluster from time-course single cell RNA sequencing experiment.**

**Supplementary Table 5: Cells identified by random forest model as corresponding to *in vivo* haematopoietic intermediates.** Numbers of cells from each timepoint which were classed as each cell type from both donors.

**Supplementary Table 6: GO terms associated with each monocle cluster from MICB^+^PVR^+^ MKP single cell RNA sequencing experiment.**

**Supplementary Table 7: Markers of each monocle cluster from MICB^+^PVR^+^ MKP single cell RNA sequencing experiment.**

**Supplementary Table 8: GSEA terms associated with each monocle cluster from MICB^+^PVR^+^ MKP single cell RNA sequencing experiment.**

## Notes

### Competing Interest Statement

The authors have declared no competing interest.

