## Supplementary figures and images for "Mapping the biogenesis of forward programmed megakaryocytes from induced pluripotent stem cells"

### Supplemental Figure 1

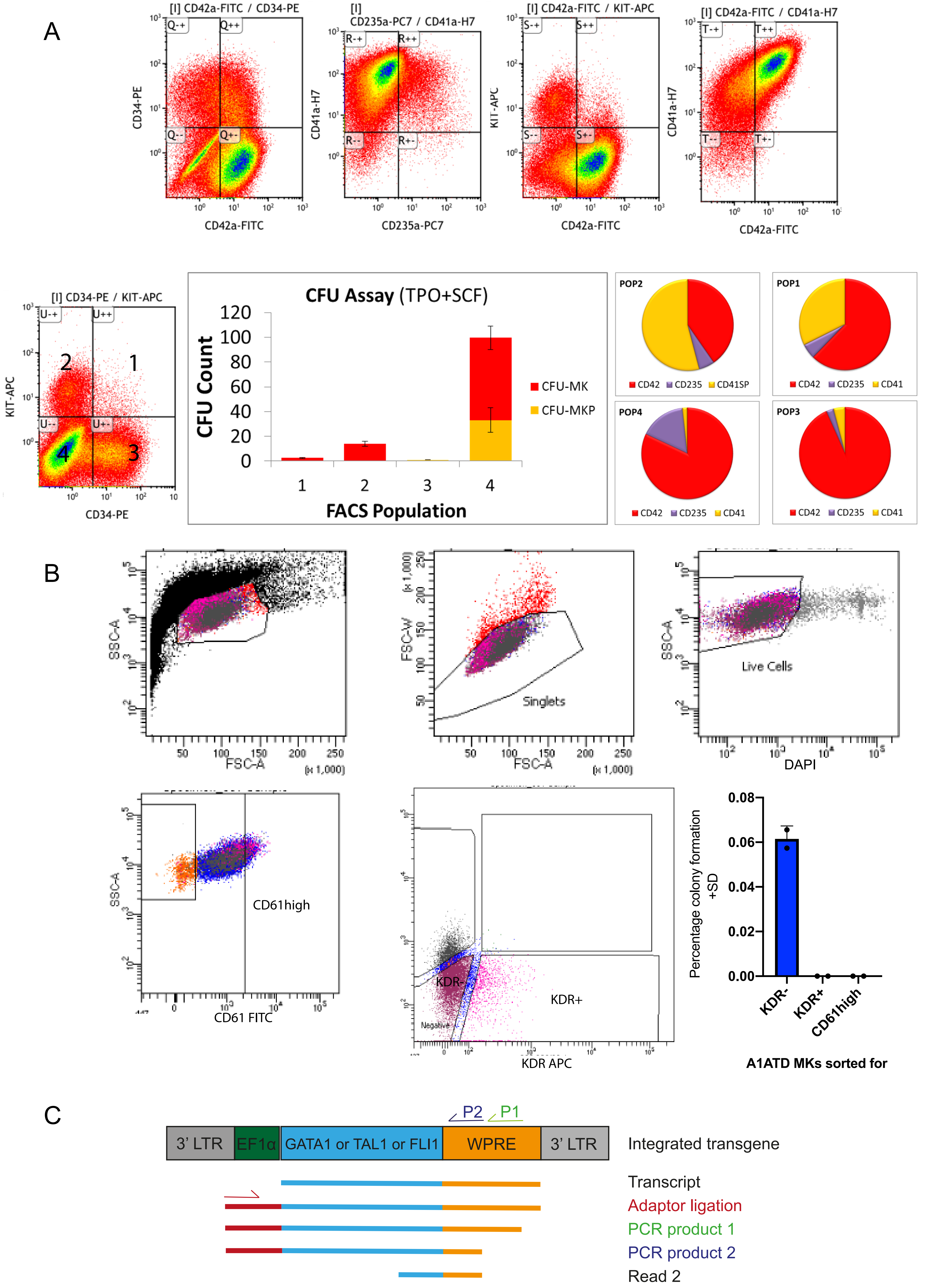

### Supplemental Figure 2

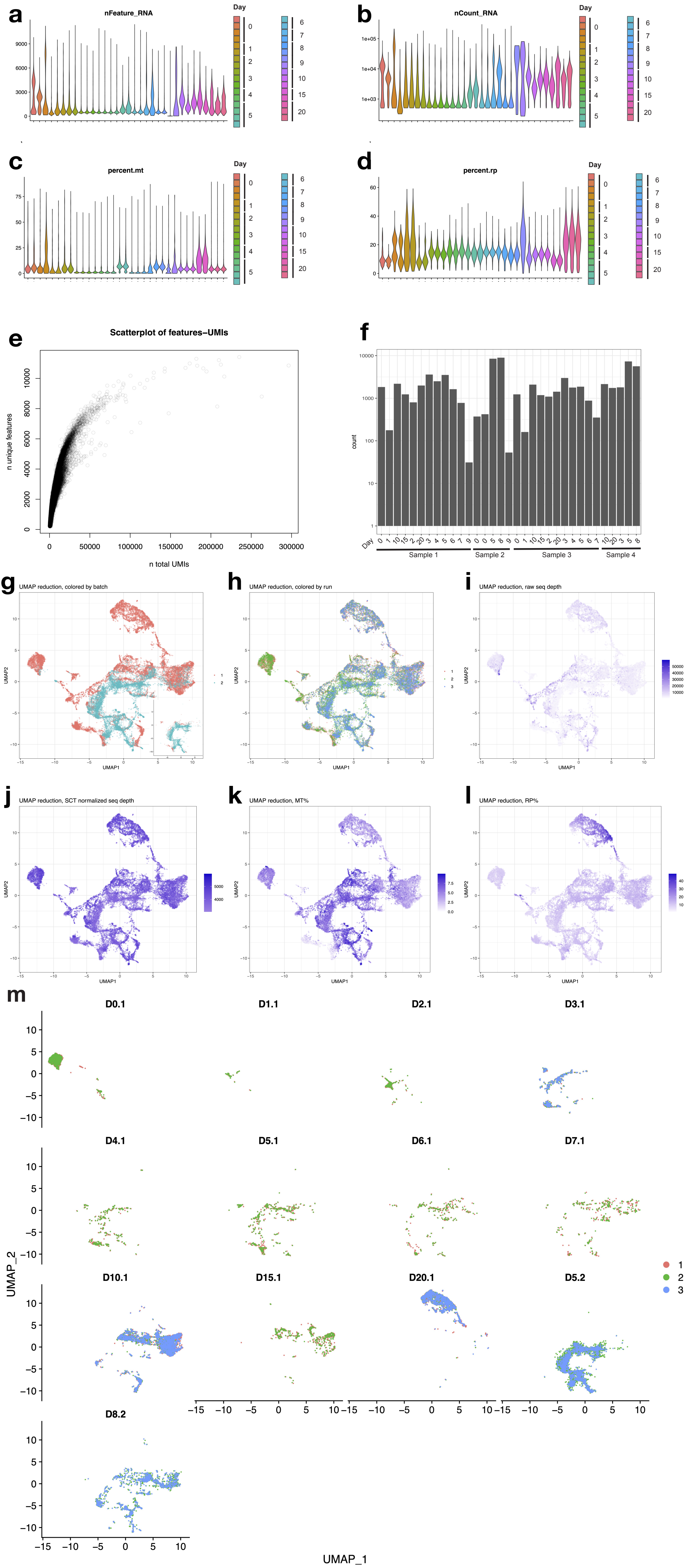

### Supplemental Figure 3

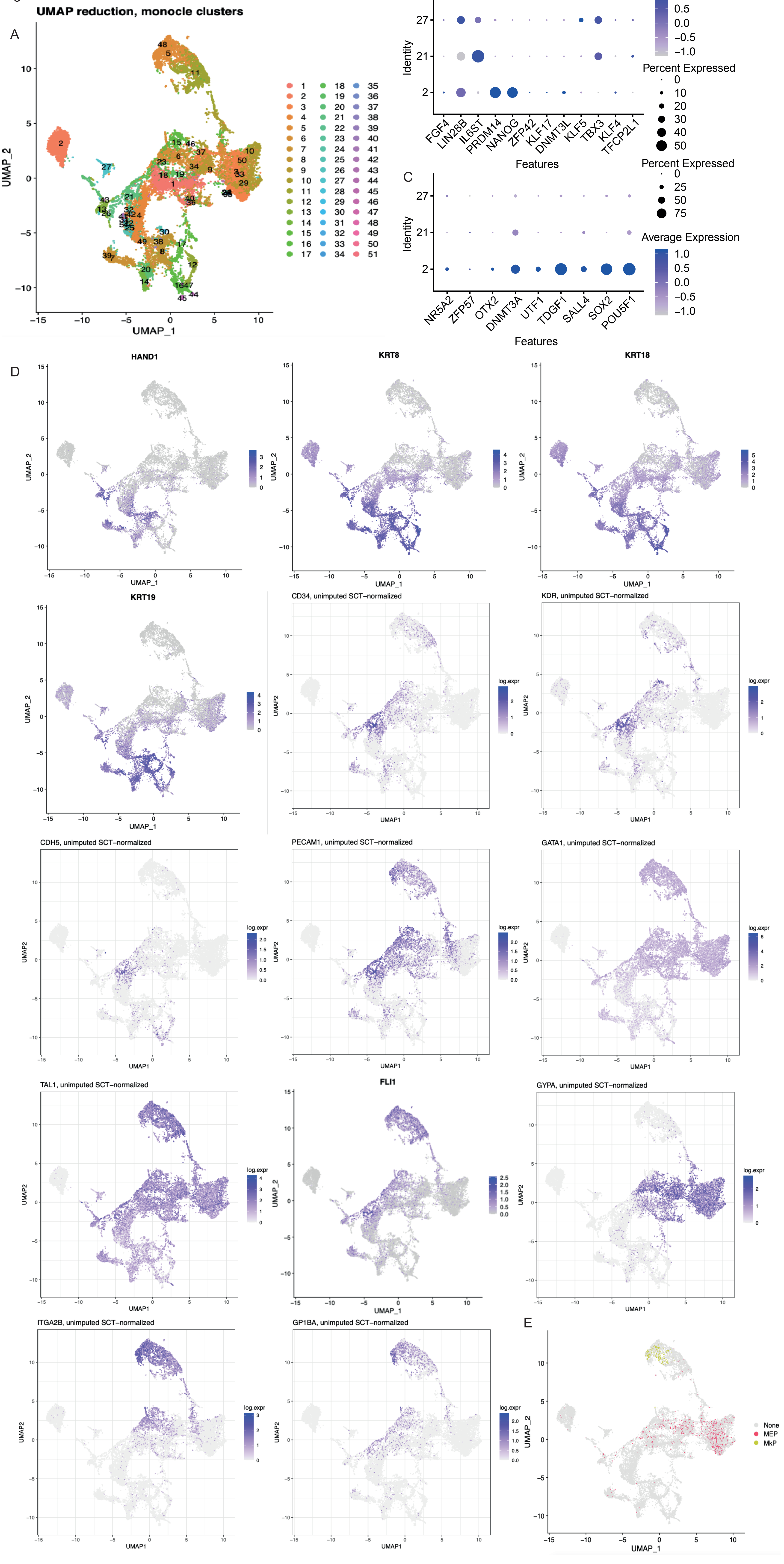

### Supplemental Figure 4

Figure S4

#### Monocle trajectory

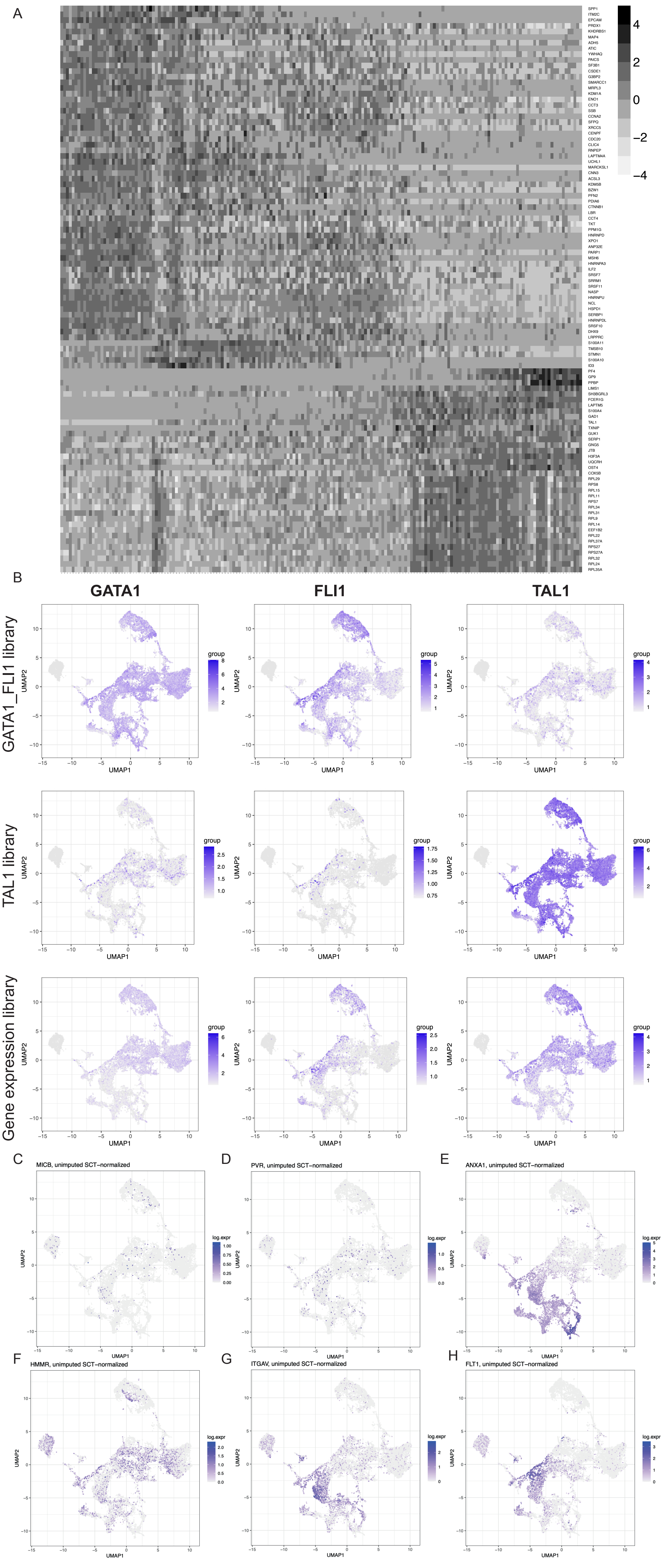

### Supplemental Figure 5

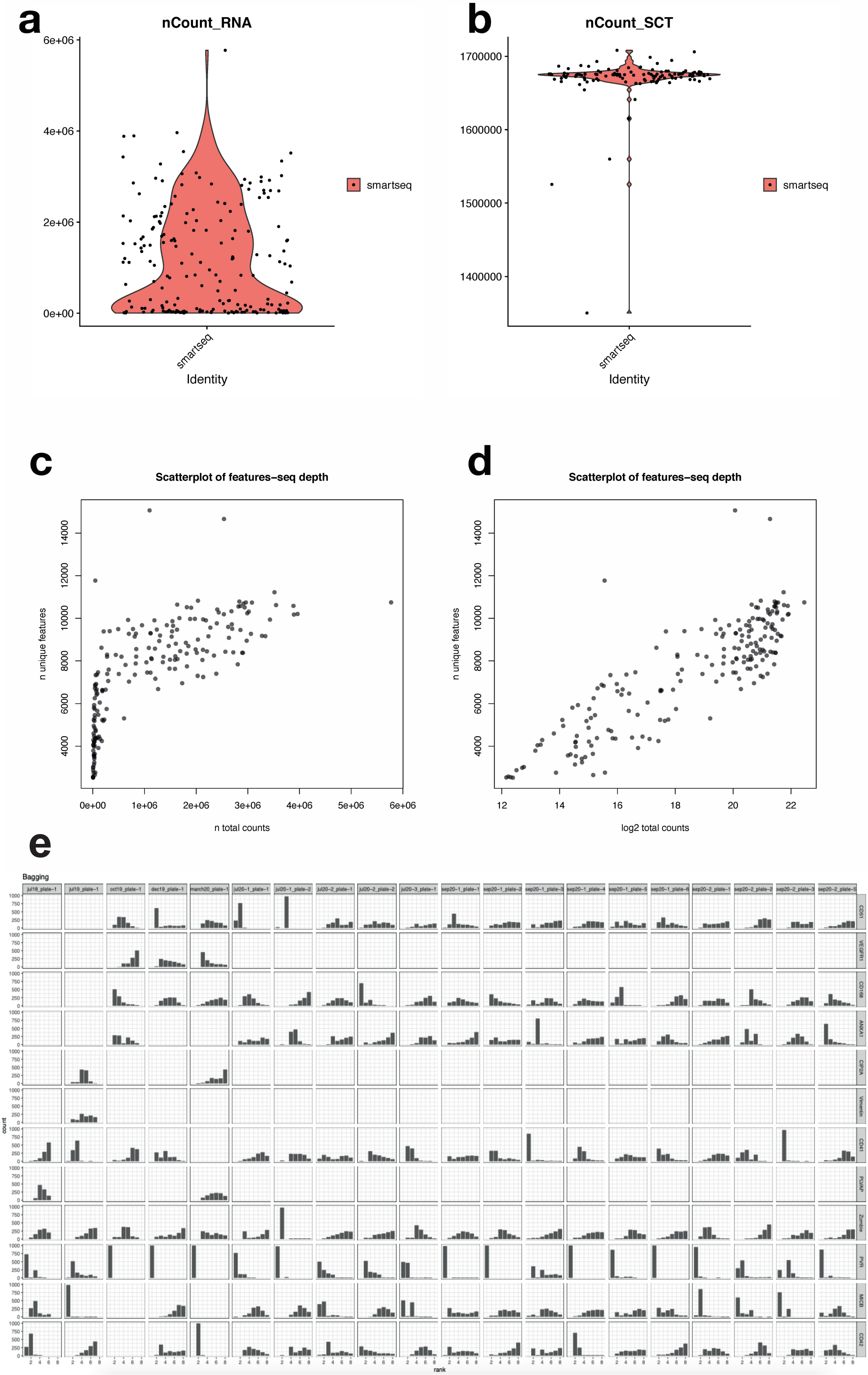

### Supplemental Figure 6

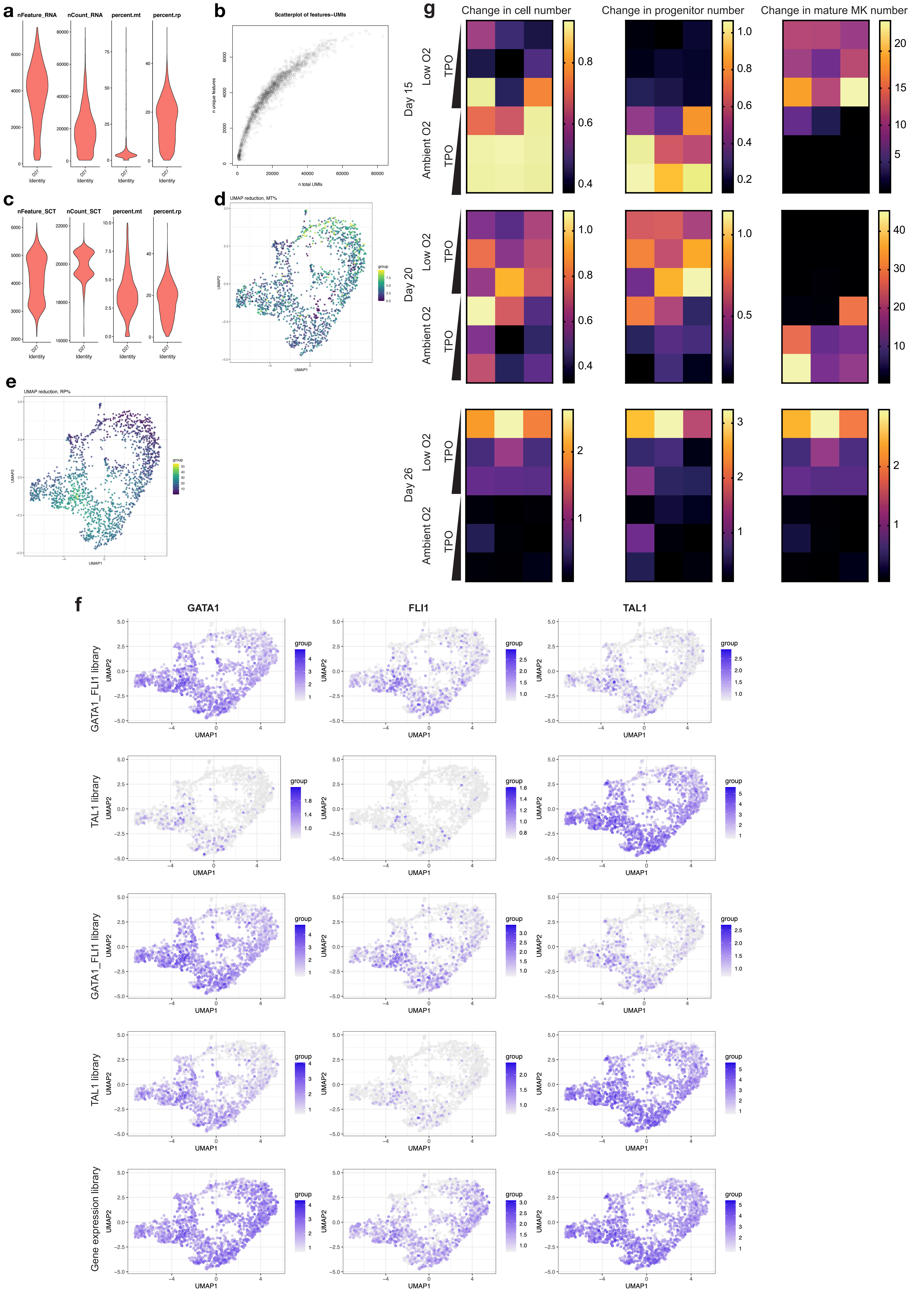
