## Supplemental Table 1 for "Mapping the biogenesis of forward programmed megakaryocytes from induced pluripotent stem cells"

**Supplementary Table 1: Primer sequences**

**qPCR primers**

| Target | Forward primer | Reverse primer |
|---|---|---|
| ITGA2B | TTCCTACTACCAGAGGCTGCATCGG | GGAGCGCCCACCAGCAGATCAT |
| GP1bA | GCCTTACACTCGCCTCACTC | TTGTGGGATAGATCCAGGGTC |
| MYB | TGCCTGGACGAACTGATAAT | GCAGGGAGTTGAGCTGTAGG |
| MICB | GCAGAGGTTCACCTGCTACA | TGGAAAGTCTGTCCGTTGACT |
| BRCA1 | AAGCAGCGGATACAACCTCA | TTCTTGATCTCCCACACTGCAA |
| CIP2A | GGAGTGGTTTGTCGGAGCAG | GGCACCAGAATAGAAAATTTTGACA |
| FEN1 | CGAACCAAGCTTTAGCCGC | AGTTTGGCCAGGCCTTGAAT |
| FANCG | AGCCTCACCCCTTCATTGTG | GGCCAGCAGGTCCAAGTAAT |
| ORC1 | GCAGCAGATCCTAAGGTCCC | GTGCATCTCCAGACAGTGCT |
| KNTC1 | GGGTACCTGAGTGTCGGTTC | GCTGCATGCCTGTATCTTTGG |
| PPIAb | GTTCTTCGACATTGCCGTCG | TCTGTGAAAGCAGGAACCCT |
| HMBS | GGGAACCAGCTCCCTGCGAAG | AGCTGTTGCCAGGATGATGGCAC |
| GAPDH | AAATCAAGTGGGGCGATGCT | CAAATGAGCCCCAGCCTTCT |
| RPL15 | GCGCCGACTGGGCTACAAGG | ACTGGGCGTTTTCGGCCACC |
| Beta globin | GCTTCTGACACAACTGTGTTCACTAGC | CACCAACTTCATCCACGTTCACC |
| Telomeric repeats | CGGTTTGTTTGGGTTTGGGTTTGGGTTTGGGTTTGGGTT | GGCTTGCCTTACCCTTACCCTTACCCTTACCCTTACCCT |

**10X amplification primers**

| Target | Sequence |
|---|---|
| GATA1 and FLI1 transgene inner | CCTCTGGATTACAAAATTTGTGAAAG |
| | GCAGCGTATCCACATAGCGT |
| | GCCATACGGGAAGCAATAGCA |
| GATA1 and FLI1 transgene outer | CACTGTGTTTGCTGACGCAAC |
| | TGCACACCACGCCAC |
| | GTGCACACCACGCCA |
| TAL1 transgene inner | CCTAGAAAAACATGGAGCAATC |
| | ATCCCAGCAGCCTAAGAACG |
| | AAGTTCAAGTCCACCGCCTT |
| TAL1 transgene outer | GGGCTAATTCACTCCCAACGA |
| | AGGCACAATCAGCATTGGTA |
| | TGTGACTGGAAAACCCACC |
