## Supplemental Table 5 for "Mapping the biogenesis of forward programmed megakaryocytes from induced pluripotent stem cells"

**Supplementary Table 5: Cells identified by random forest model as corresponding to *in vivo* haematopoietic intermediates.**

| Donor 1 | EryP | HSC/MPP | MEP | MkP | None |
|---|---|---|---|---|---|
| D0 | 0 | 0 | 11 | 0 | 1557 |
| D0_single | 0 | 0 | 0 | 0 | 103 |
| D0_timecour | 0 | 0 | 0 | 0 | 36 |
| D1 | 0 | 0 | 3 | 0 | 98 |
| D2 | 0 | 0 | 4 | 0 | 448 |
| D3 | 0 | 1 | 1 | 0 | 1155 |
| D4 | 0 | 0 | 6 | 0 | 460 |
| D5 | 8 | 42 | 121 | 0 | 5986 |
| D6 | 5 | 4 | 20 | 0 | 423 |
| D7 | 12 | 6 | 36 | 0 | 390 |
| D8 | 26 | 38 | 262 | 1 | 2077 |
| D9 | 2 | 0 | 13 | 0 | 30 |
| D10 | 49 | 13 | 1892 | 0 | 2984 |
| D15 | 6 | 0 | 265 | 25 | 436 |
| D20 | 0 | 0 | 341 | 964 | 1876 |

| Donor 2 | | | MEP | MkP | None |
|---|---|---|---|---|---|
| D0 | | | 15 | 0 | 1553 |
| D0_single | | | 2 | 0 | 101 |
| D0_timecourse | | | 0 | 0 | 36 |
| D1 | | | 3 | 0 | 98 |
| D2 | | | 9 | 0 | 443 |
| D3 | | | 28 | 0 | 1129 |
| D4 | | | 18 | 0 | 448 |
| D5 | | | 64 | 0 | 6093 |
| D6 | | | 11 | 0 | 441 |
| D7 | | | 26 | 0 | 418 |
| D8 | | | 100 | 0 | 2304 |
| D9 | | | 0 | 0 | 45 |
| D10 | | | 556 | 0 | 4382 |
| D15 | | | 43 | 3 | 686 |
| D20 | | | 26 | 289 | 2866 |
