## Supplemental Table 7 for "Mapping the biogenesis of forward programmed megakaryocytes from induced pluripotent stem cells"

**Supplementary Table 7: Markers of each monocle cluster from MICB+PVR+ MKP scRNA seq**

| avg_diff | power | avg_logFC | pct.1 | pct.2 | cluster | gene |
|---|---|---|---|---|---|---|
| 1.89439945 | 0.802 | 1.89439945 | 0.926 | 0.453 | 1 | S100A11 |
| 1.06680347 | 0.784 | 1.06680347 | 1 | 0.955 | 1 | RAC2 |
| 1.17547787 | 0.742 | 1.17547787 | 0.953 | 0.631 | 1 | SAMSN1 |
| 1.21874457 | 0.738 | 1.21874457 | 1 | 0.975 | 1 | PRDX1 |
| 1.72911231 | 0.722 | 1.72911231 | 0.973 | 0.755 | 1 | VIM |
| 1.26858477 | 0.704 | 1.26858477 | 0.791 | 0.145 | 1 | NAV1 |
| 1.5164418 | 0.7 | 1.5164418 | 0.899 | 0.71 | 1 | VWA5A |
| 1.9891221 | 0.67 | 1.9891221 | 0.716 | 0.088 | 1 | ANXA1 |
| 1.07106465 | 0.668 | 1.07106465 | 0.743 | 0.122 | 1 | ARHGEF6 |
| 1.22294348 | 0.646 | 1.22294348 | 0.811 | 0.209 | 1 | GDF3 |
| 3.0943309 | 0.63 | 3.0943309 | 0.642 | 0.022 | 1 | CPA3 |
| 1.58294622 | 0.61 | 1.58294622 | 0.628 | 0.027 | 1 | AHNAK |
| 1.13182834 | 0.61 | 1.13182834 | 0.831 | 0.496 | 1 | CD99 |
| 2.18253141 | 0.604 | 2.18253141 | 0.622 | 0.042 | 1 | KIT |
| 1.84345316 | 0.604 | 1.84345316 | 0.608 | 0.007 | 1 | HPGDS |
| 1.10409351 | 0.594 | 1.10409351 | 0.892 | 0.463 | 1 | BST2 |
| 1.039758 | 0.574 | 1.039758 | 0.669 | 0.149 | 1 | PMP22 |
| 1.22820618 | 0.536 | 1.22820618 | 0.831 | 0.597 | 1 | SFRP1 |
| 2.12903661 | 0.528 | 2.12903661 | 0.561 | 0.043 | 1 | NTS |
| 1.99817627 | 0.522 | 1.99817627 | 0.534 | 0.017 | 1 | RHEX |
| 1.02154036 | 0.854 | 1.02154036 | 0.994 | 0.861 | 2 | RANBP1 |
| 1.24584739 | 0.816 | 1.24584739 | 0.99 | 0.867 | 2 | DUT |
| 1.42631717 | 0.808 | 1.42631717 | 0.9 | 0.213 | 2 | FABP5 |
| 1.8860931 | 0.804 | 1.8860931 | 0.881 | 0.204 | 2 | TYMS |
| 1.40129913 | 0.8 | 1.40129913 | 1 | 0.979 | 2 | TUBA1B |
| 1.6171925 | 0.796 | 1.6171925 | 0.942 | 0.488 | 2 | UBE2T |
| 1.01885096 | 0.788 | 1.01885096 | 0.984 | 0.549 | 2 | SRM |
| 1.07889792 | 0.786 | 1.07889792 | 0.99 | 0.756 | 2 | LDHA |
| 1.28752423 | 0.734 | 1.28752423 | 0.842 | 0.229 | 2 | MCM7 |
| 1.3064557 | 0.716 | 1.3064557 | 0.791 | 0.139 | 2 | PCLAF |
| 1.42193273 | 0.698 | 1.42193273 | 0.945 | 0.633 | 2 | STMN1 |
| 1.02385994 | 0.698 | 1.02385994 | 0.826 | 0.242 | 2 | MCM3 |
| 1.09408238 | 0.694 | 1.09408238 | 0.814 | 0.186 | 2 | MCM5 |
| 1.03756463 | 0.678 | 1.03756463 | 0.875 | 0.37 | 2 | MGST1 |
| 1.07438355 | 0.67 | 1.07438355 | 0.73 | 0.096 | 2 | ZWINT |
| 1.01859475 | 0.67 | 1.01859475 | 0.727 | 0.095 | 2 | ATAD2 |
| 1.03291006 | 0.666 | 1.03291006 | 0.746 | 0.104 | 2 | CDT1 |
| 1.04454366 | 0.66 | 1.04454366 | 0.849 | 0.367 | 2 | PCNA |
| 1.01334835 | 0.644 | 1.01334835 | 0.74 | 0.147 | 2 | FEN1 |
| 1.10211838 | 0.634 | 1.10211838 | 0.707 | 0.122 | 2 | CENPW |
| 1.0727987 | 0.824 | 1.0727987 | 1 | 1 | 3 | OAZ1 |
| 1.01065223 | 0.778 | 1.01065223 | 0.997 | 0.85 | 3 | P2RX1 |
| 1.10698415 | 0.77 | 1.10698415 | 0.992 | 0.853 | 3 | MFSD1 |
| 2.02462446 | 0.764 | 2.02462446 | 0.97 | 0.55 | 3 | CLEC1B |

| | | | | | | |
|---|---|---|---|---|---|---|
| 1.03131684 | 0.76 | 1.03131684 | 0.985 | 0.819 | 3 | NT5C3A |
| 2.44404412 | 0.754 | 2.44404412 | 0.955 | 0.575 | 3 | PPBP |
| 1.46813494 | 0.744 | 1.46813494 | 0.99 | 0.731 | 3 | TUBB1 |
| 1.14055032 | 0.73 | 1.14055032 | 0.985 | 0.749 | 3 | TMEM40 |
| 1.14883427 | 0.724 | 1.14883427 | 0.965 | 0.718 | 3 | CNST |
| 1.17528564 | 0.718 | 1.17528564 | 0.945 | 0.584 | 3 | DMTN |
| 1.06183384 | 0.714 | 1.06183384 | 1 | 0.971 | 3 | STOM |
| 1.11400315 | 0.712 | 1.11400315 | 0.997 | 0.858 | 3 | CTSA |
| 1.07282427 | 0.706 | 1.07282427 | 0.975 | 0.762 | 3 | ADD3 |
| 1.27138942 | 0.704 | 1.27138942 | 0.987 | 0.772 | 3 | TREML1 |
| 1.09121325 | 0.67 | 1.09121325 | 0.992 | 0.844 | 3 | GP9 |
| 1.00979604 | 0.668 | 1.00979604 | 0.977 | 0.67 | 3 | LTBP1 |
| 1.33767459 | 0.656 | 1.33767459 | 0.982 | 0.786 | 3 | THBS1 |
| 1.1422708 | 0.648 | 1.1422708 | 0.977 | 0.726 | 3 | GSN |
| 1.4494833 | 0.646 | 1.4494833 | 0.972 | 0.734 | 3 | TUBA4A |
| 1.0705744 | 0.646 | 1.0705744 | 0.97 | 0.731 | 3 | ABCC3 |
| 1.02628691 | 0.882 | 1.02628691 | 1 | 0.986 | 4 | PSMB2 |
| 1.04517959 | 0.824 | 1.04517959 | 1 | 0.995 | 4 | HSP90AB1 |
| 1.00886839 | 0.768 | 1.00886839 | 0.991 | 0.853 | 4 | CYB5R1 |
| 1.6718063 | 0.722 | 1.6718063 | 0.817 | 0.217 | 4 | MLLT11 |
| 1.0993491 | 0.64 | 1.0993491 | 1 | 0.964 | 4 | SQSTM1 |
| 2.08337355 | 0.566 | 2.08337355 | 0.948 | 0.847 | 4 | HSPA1A |
| 1.23826808 | 0.502 | 1.23826808 | 1 | 0.998 | 4 | HSP90AA1 |
| 1.19127403 | 0.44 | 1.19127403 | 0.983 | 0.931 | 4 | HSPA5 |
| 1.13492704 | 0.418 | 1.13492704 | 0.948 | 0.803 | 4 | SLC3A2 |
| 1.18768431 | 0.408 | 1.18768431 | 0.426 | 0.021 | 4 | UCHL1 |
| 2.14999504 | 0.906 | 2.14999504 | 0.989 | 0.445 | 5 | ALAS2 |
| 1.79723561 | 0.84 | 1.79723561 | 0.966 | 0.604 | 5 | GYPC |
| 1.46968829 | 0.82 | 1.46968829 | 1 | 0.987 | 5 | BLVRB |
| 2.45621281 | 0.816 | 2.45621281 | 0.862 | 0.18 | 5 | HMBS |
| 1.61960671 | 0.804 | 1.61960671 | 0.966 | 0.518 | 5 | AC104389.4 |
| 1.51993942 | 0.782 | 1.51993942 | 0.954 | 0.548 | 5 | BTG2 |
| 2.36113983 | 0.766 | 2.36113983 | 0.931 | 0.503 | 5 | HBA2 |
| 1.70911405 | 0.766 | 1.70911405 | 0.828 | 0.12 | 5 | GYPA |
| 1.68743111 | 0.758 | 1.68743111 | 0.885 | 0.306 | 5 | GYPB |
| 1.96806172 | 0.746 | 1.96806172 | 0.977 | 0.828 | 5 | SLC25A37 |
| 1.73604509 | 0.734 | 1.73604509 | 0.954 | 0.784 | 5 | HBA1 |
| 1.60580761 | 0.72 | 1.60580761 | 0.989 | 0.886 | 5 | HBG2 |
| 1.4689631 | 0.696 | 1.4689631 | 0.839 | 0.273 | 5 | KLF1 |
| 1.43956146 | 0.696 | 1.43956146 | 0.897 | 0.503 | 5 | KRT18 |
| 1.44912709 | 0.682 | 1.44912709 | 0.828 | 0.29 | 5 | CTSL |
| 1.02911898 | 0.662 | 1.02911898 | 0.977 | 0.74 | 5 | FAM210B |
| 3.83662041 | 0.654 | 3.83662041 | 0.828 | 0.334 | 5 | HBE1 |
| 2.33381902 | 0.62 | 2.33381902 | 0.966 | 0.847 | 5 | HBG1 |
| 1.61500841 | 0.6 | 1.61500841 | 0.897 | 0.522 | 5 | CTSB |
| 4.28965627 | 0.594 | 4.28965627 | 0.655 | 0.116 | 5 | HBZ |

| | | | | | | |
|---|---|---|---|---|---|---|
| 1.18443835 | 0.594 | 1.18443835 | 0.966 | 0.829 | 5 | SLC25A39 |
| 1.47222499 | 0.79 | 1.47222499 | 0.92 | 0.231 | 7 | GDF3 |
| 1.3629003 | 0.764 | 1.3629003 | 0.973 | 0.479 | 7 | BST2 |
| 1.47917382 | 0.76 | 1.47917382 | 0.973 | 0.765 | 7 | VIM |
| 1.05482845 | 0.726 | 1.05482845 | 0.92 | 0.371 | 7 | CD44 |
| 1.41423431 | 0.714 | 1.41423431 | 0.84 | 0.259 | 7 | ECSCR |
| 1.0276979 | 0.676 | 1.0276979 | 0.987 | 0.977 | 7 | PRDX1 |
| 1.38916534 | 0.662 | 1.38916534 | 0.8 | 0.294 | 7 | ENC1 |
| 1.34550534 | 0.608 | 1.34550534 | 0.92 | 0.68 | 7 | IGFBP2 |
| 1.04236713 | 0.608 | 1.04236713 | 0.747 | 0.186 | 7 | ALDH1A1 |
| 1.09692829 | 0.606 | 1.09692829 | 1 | 0.988 | 7 | DBI |
| 1.016274 | 0.564 | 1.016274 | 0.72 | 0.206 | 7 | PLVAP |
| 1.02064845 | 0.544 | 1.02064845 | 0.76 | 0.232 | 7 | TM4SF1 |
| 1.51825271 | 0.896 | 1.51825271 | 1 | 0.993 | 9 | RPS27L |
| 2.02508285 | 0.798 | 2.02508285 | 1 | 0.666 | 9 | CDKN1A |
| 1.21854035 | 0.744 | 1.21854035 | 0.983 | 0.555 | 9 | BTG2 |
| 1.33760033 | 0.742 | 1.33760033 | 0.966 | 0.616 | 9 | BBC3 |
| 1.19493358 | 0.716 | 1.19493358 | 0.845 | 0.236 | 9 | PIK3IP1 |
| 1.31237186 | 0.7 | 1.31237186 | 1 | 0.883 | 9 | UBE2B |
| 1.19234282 | 0.692 | 1.19234282 | 1 | 0.873 | 9 | HIST1H1C |
| 1.23904757 | 0.678 | 1.23904757 | 0.828 | 0.375 | 9 | ANXA4 |
| 1.0058897 | 0.674 | 1.0058897 | 1 | 0.978 | 9 | NEAT1 |
| 1.84998797 | 0.67 | 1.84998797 | 1 | 0.861 | 9 | SAT1 |
| 1.05959657 | 0.67 | 1.05959657 | 1 | 0.908 | 9 | ISCU |
| 1.90597088 | 0.654 | 1.90597088 | 0.879 | 0.532 | 9 | GDF15 |
| 1.08214888 | 0.654 | 1.08214888 | 1 | 0.799 | 9 | HIST1H2AC |
| 1.06212328 | 0.654 | 1.06212328 | 0.897 | 0.574 | 9 | STX7 |
| 1.34080439 | 0.626 | 1.34080439 | 0.862 | 0.498 | 9 | DDIT3 |
| 1.30809635 | 0.612 | 1.30809635 | 0.983 | 0.819 | 9 | IFIT1 |
| 1.09568568 | 0.61 | 1.09568568 | 0.845 | 0.535 | 9 | TRIAP1 |
| 1.13203514 | 0.58 | 1.13203514 | 0.879 | 0.548 | 9 | BTG1 |
| 1.79819706 | 0.576 | 1.79819706 | 0.672 | 0.192 | 9 | HIST1H4H |
| 1.08078227 | 0.57 | 1.08078227 | 0.707 | 0.184 | 9 | TMEM140 |
| 1.05839548 | 0.456 | 1.05839548 | 0.893 | 0.685 | 11 | SH3BP5 |
| 1.14985343 | 0.92 | 1.14985343 | 1 | 0.967 | 13 | RBM39 |
| 1.49857256 | 0.834 | 1.49857256 | 1 | 0.786 | 13 | RSRP1 |
| 2.18087255 | 0.8 | 2.18087255 | 1 | 0.998 | 13 | MALAT1 |
| 1.93877545 | 0.726 | 1.93877545 | 1 | 0.979 | 13 | NEAT1 |
| 1.0648091 | 0.692 | 1.0648091 | 1 | 0.689 | 13 | AFF4 |
| 1.01956665 | 0.648 | 1.01956665 | 1 | 0.788 | 13 | N4BP2L2 |
| 1.00064124 | 0.634 | 1.00064124 | 0.944 | 0.843 | 13 | WSB1 |
| 1.02926201 | 0.63 | 1.02926201 | 0.944 | 0.946 | 13 | LUC7L3 |
| 1.00863521 | 0.466 | 1.00863521 | 1 | 0.927 | 13 | PPP1R15A |
| 1.18355185 | 0.43 | 1.18355185 | 0.889 | 0.551 | 13 | SERTAD1 |
